# Combined nanometric and phylogenetic analysis of unique endocytic compartments in Giardia lamblia sheds light on the evolution of endocytosis in Fornicata

**DOI:** 10.1101/2022.04.14.488357

**Authors:** Rui Santos, Ásgeir Ástvaldsson, Shweta V. Pipaliya, Jon Paulin Zumthor, Joel B. Dacks, Staffan Svärd, Adrian B. Hehl, Carmen Faso

## Abstract

*Giardia lamblia*, a parasitic protist of the metamonada supergroup, has evolved one of the most diverged endocytic compartment systems investigated so far. Peripheral endocytic compartments, currently known as peripheral vesicles or vacuoles (PVs), perform bulk uptake of fluid phase material which is then digested and sorted either to the cell cytosol or back to the extracellular space. Here, we present a quantitative morphological characterization of these organelles using volumetric electron microscopy and super-resolution microscopy (SRM). We defined a morphological classification for the heterogenous population of PVs and performed a comparative analysis of PVs and endosome-like organelles in representatives of phylogenetically-related taxa, *Spironucleus spp.* and *Tritrichomonas foetus*. To investigate the as-yet insufficiently understood connection between PVs and clathrin assemblies in *G. lamblia*, we further performed an in-depth search for two key elements of the endocytic machinery, clathrin heavy chain (CHC) and clathrin light chain (CLC) across different lineages in Metamonada. Our data point to the loss of a *bona fide* CLC in the last Fornicata common ancestor (LFCA) with the emergence of a protein analogous to CLC (*Gl*ACLC) in the *Giardia* genus. Taken together, this provides the first comprehensive nanometric view of *Giardia*’s endocytic system architecture and sheds light on the evolution of GLACLC analogues in the Fornicata supergroup and, specific to Giardia, as a possible adaptation to the formation and maintenance of stable clathrin assemblies at PVs.

## INTRODUCTION

Endomembrane compartments, while present in a few prokaryotic lineages (Heimerl et al. 2017), have evolved and greatly diversified across eukaryotic lineages. A fundamental task performed by some membrane-bounded organelles is endocytosis – the controlled and directed uptake of nutrients and other materials from the extracellular space into the cell by membrane transport. Fluid phase or receptor-bound material at the cell surface is internalised via invaginations and formation of vesicles at the plasma membrane, mediated by clathrin-coated vesicles (CCVs) (Robinson 2015; Kaksonen and Roux 2018). In turn, CCVs fuse with early endosomes which mature into late endosomes upon lysosome fusion (Huotari and Helenius 2011; Naslavsky and Caplan 2018). Clathrin coats are also involved in protein secretion forming exocytic transport vesicles derived from the trans-Golgi compartment and play a role in Golgi apparatus reassembly after mitotic cell division (Radulescu et al. 2007; Jaiswal et al. 2009).

Evolutionary adaptations of endocytic pathways to specific environmental niches and nutrient sources are especially relevant to species adopting a fully parasitic or commensal lifestyle (Poulin and Randhawa 2015; Jackson et al. 2016; Dacks and Field 2018; Pipaliya et al. 2021).

Within the extant Metamonada supergroup (Hampl et al. 2009; Hug et al. 2016; Burki et al. 2020), the parasitic protist *Giardia lamblia* (syn.: *intestinalis* or *duodenalis*) evolved a distinct endocytic pathway, which reflects its adaptation to the host intestinal lumen environment. This unicellular parasite is responsible for >300 million cases annually of water-borne infections causing gastroenteritis – giardiasis – with higher incidence in low to middle income countries (Caccìo and Ryan 2008) (Caccìo and Ryan 2008). *Giardia* is the etiological agent for symptomatic gastroenteritis in 15% of children in developing countries, with 1-2% fatality in children with severely compromised health status (Kotloff et al. 2013; Lanata et al. 2013). There is a strong association of *Giardia* infections with chronic conditions such as irritable bowel syndrome or inflammatory bowel disease as a result of intestinal barrier function disruption and microbiome dysregulation (Allain et al. 2017; Fekete et al. 2021).

Cellular evolution of the *Giardia* genus as an obligate parasite adapted to the small intestinal niche of vertebrates is characterised by a reduction in subcellular compartment diversity. Peroxisomes, late endosomes and a permanent stacked Golgi complex have not been detected in *Giardia* (Faso and Hehl 2011). Two nuclei (Benchimol 2005), an extensive endoplasmic reticulum (ER) (Soltys et al. 1996), highly reduced mitochondria-derived organelles – the mitosomes (Tovar et al. 2003) - and peripheral vesicles (PVs) (Lanfredi-Rangel et al. 1998) are the only membrane-bounded organelles with conserved morphology and function documented in the *Giardia* trophozoite (Marti, Regös, et al. 2003; Zumthor et al. 2016; Cernikova et al. 2020).

The complex array of PV organelles as the only documented endocytic membrane compartment system in *Giardia* is responsible for uptake of fluid-phase and membrane-bound material (Rivero et al. 2011; Zumthor et al. 2016; Frontera et al. 2018). These organelles acidify and presumably serve as digestive compartments with capability for sorting after processing, similar to early and late endosomes and lysosomes (Lanfredi-Rangel et al. 1998). The static system of PV organelles (Abodeely et al. 2009; Zumthor et al. 2016) is restricted to the peripheral cortex below the plasma membrane (PM) of the *Giardia* trophozoite. PV morphology was investigated using high-resolution electron microscopy serial sectioning and three-dimensional reconstruction (Zumthor et al. 2016). These organelles were resolved as tubular structures in close proximity to funnel-shaped invaginations of the PM (Zumthor et al. 2016). In the same report, the presence of focal accumulations of clathrin heavy chain (CHC) molecules and their main interactors, collectively termed clathrin assemblies, were demonstrated at PM and PV membrane interfaces. The function of these stable focal assemblies as well as additional components at the interface of the PV membranes and the PM, has proved elusive (Zumthor et al. 2016). However, transient association of several members of the family of adaptor proteins (AP) suggests a role in dynamic processes linked to uptake of fluid-phase and receptor-bound material into PVs (Zumthor et al. 2016; Cernikova et al. 2020). Our current working model for bulk fluid-phase uptake of extracellular material into PVs invokes a “kiss and flush” mechanism, whereby acidified PV membranes and the PM transiently form channels at invaginations allowing exchange between PV lumen content and the extracellular space at regular intervals. Endocytosed material is digested in the sealed-off acidified PVs and transported towards the cell interior while residual material and waste is flushed to the extracellular space in the next round of membrane fusion, thus completing the PV cycle (Zumthor et al. 2016; Cernikova et al. 2020).

In this report, we address open questions concerning *G. lamblia’s* PV ultrastructure and its associated molecular machinery in a comparative approach with one closely and one more distantly related fornicata and metamonada species, *Spironucleus sp*. and *Tritrichomonas foetu*s, respectively. Using volumetric electron microscopy and super resolution light microscopy we developed a classification of PVs based on organelle morphology. Comparative analysis of *Giardia’s* PVs with endocytic compartments of fornicata and metamonada species emphasized the genus-specific nature of the *Giardia* endocytic system architecture. In addition, using a combination of co-immunoprecipitation and phylogeny techniques, we provide evidence that a proposed diverged clathrin light chain previously named *GlCLC* (Zumthor et al. 2016) is unique to the *Giardia* genus and evolved *de novo* as structurally analogous to CLC after loss of a *bona fide* CLC in the last Fornicata common ancestor (LFCA). Taken together, the emergence of a unique and highly polymorphic endocytic system such as the one found in the genus *Giardia* is linked to the proposed convergent evolution of an independent CLC analogue concomitant with loss of a mostly conserved CLC orthologue.

## RESULTS AND DISCUSSION

### Complete FIB-SEM rendering of a *G. lamblia* trophozoite reveals a novel landscape of vesicular compartments

Volumetric scanning electron microscopy (vSEM) is currently considered the gold standard for the determination of biological ultrastructure (Titze and Genoud 2016). Focused ion beam electron scanning microscopy (FIB-SEM) uses a beam of gallium ions to mill and image consecutive layers of an embedded biological sample, resulting in a voxel resolution as low as 1-2 nm (Kizilyaprak et al. 2014). This technique allows for sectioning and imaging of entire cells (Wei et al. 2012). It was previously implemented for partial rendering of *G. lamblia* trophozoite sections (Zumthor et al. 2016) and more recently in (Tůmová et al. 2020). Here we sectioned for the first time a complete *G. lamblia* trophozoite at a voxel resolution of 125 nm^3^ (5x5x5 nm) after high pressure freezing (HPF) and embedding. Images representing the sagittal plane adjacent to the cell centre (figure 1A, 1D and supplementary figure 1A and B) show all the major cell compartments such as the nuclei (Figure 1A, N), the endoplasmic reticulum (Figure 1A, ER), mitosomes (Figure 1A and C, m) and elements of the cytoskeleton: axonemes (Ax), funis (F) and the ventral disc (VD) (Figure 1D; Dawson 2010). Two different types of small cytoplasmic organelles are observed: PVs (arrow heads) with heterogenous morphology and smaller and electron-dense membrane vesicles of uniform size and appearance which we termed Small Vesicles (SVs; asterisks) (figure 1B and E).

**Figure 1.**
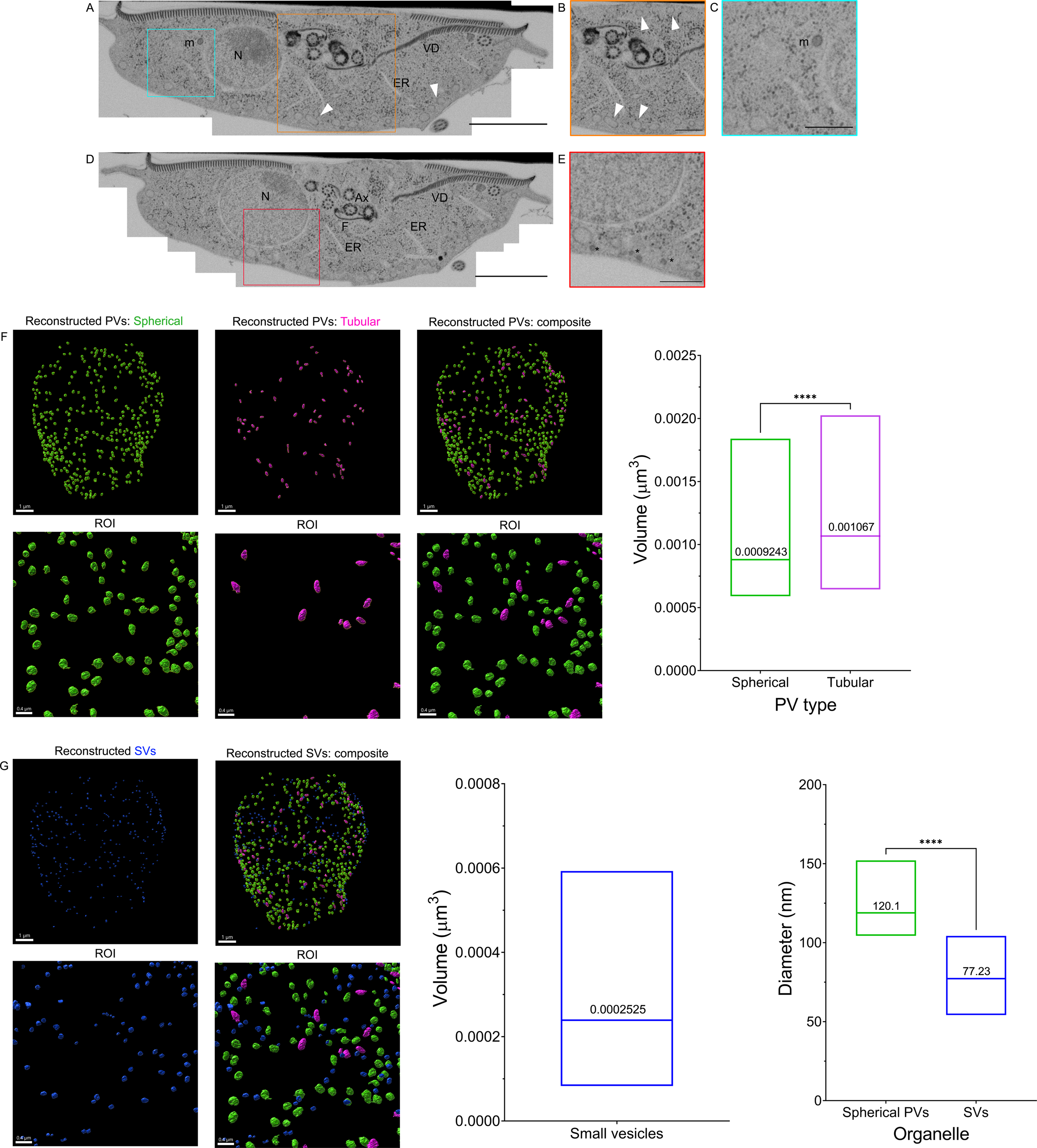
Complete scanning of a *Giardia* trophozoite by focused ion beam scanning electron microscopy (FIB-SEM). (A and D) A complete *G. lamblia* trophozoite was scanned with an isotropic resolution of 5 nm. The Nucleus (N), the Endoplasmic Reticulum (ER), cytoskeletal features such as the Ventral disk (VD), Funis (F) and axonemes (Ax) and mitosomes (m), are observed. Peripheral vesicles (PV) are marked by arrowheads. Smaller, electron dense vesicles are also documented (asterisks) – small vesicles (SVs). (B) Region of interest of (A) highlighting PVs of different morphology with arrowheads. (C) Highlight of mitosomes proximal to ER membrane. (E) Region of interest of (D) highlighting SVs (asterisks). (F) Full reconstruction of PVs with ilastik and rendering in Imaris reveals the presence of at least two PV morphologies: spherical (green) and tubular (violet). Box-plot: 403 spherical and 64 tubular PVs were segmented out. Spherical PVs average a volume of 0.0009243±0.0003322 μm^3^ with a 95% confidence interval between [0.0009022; 0.0009022] μm3. Tubular PVs average a volume of 0.001067±0.0003322μm^3^ with a 95% confidence interval between [0.0009843; 0.001150] μm^3^. (G) Reconstruction of 269 SVs revealed their volumes average 0.0002525±9.280x10^-5^ μm^3^ in a 95% confidence interval between [0.0002414; 0.0002637] μm^3^ (left box-plot). This equals to an average diameter of 77.23±9.666 nm in a 95% confidence interval between [76.07; 78.40] nm. The diameter of spherical PVs averages 120.1±9.507 nm in a 95% confidence interval between [119.2; 121] nm (right box-plot). The difference in diameter between SVs and spherical PVs is statistically significant (****; p-value < 0.0001, t-student significance test). Scale bars: (A and D) 2 μm, (B, C and E) 500 nm. ROI: region of interest.

After serial sectioning and alignment with TrakEM (Cardona et al. 2012), we used the supervised machine learning (ML) tool Ilastik for pixel based image segmentation of PVs and SVs (Sommer et al. 2011; Berg et al. 2019). The algorithm collection performs supervised learning and recognition of patterns based on ground truth training provided by the user. Patterns are sorted into classes. Once the algorithm is trained on a subset of image data, it is used to analyse complete datasets and assigning features to different classes following a decision tree method (Sommer and Gerlich 2013; Kan 2017). This process enabled the three-dimensional rendering of selected trophozoite features: the complete cytoskeleton, the ER, PVs and mitosomes (supplementary figure 1C). In addition, we were able to calculate the volume of the cell at 138 μm^3^ as well as the average volume of mitosome organelles (N=14) at 0.001093±0.0005698μm^3^ with a 95% confidence interval between [0.0007643, 0.001422] μm^3^ (supplementary figure 1D).

Similarly, supervised ML assisted pixel segmentation and object clustering analysis allowed identification of two statistically distinct morphological classes of PVs: spherical and tubular/elongated PVs. Individual PV organelles of both classes (N=467) were rendered in three dimensions (figure 1F and supplementary video 1). Spherical PVs average a volume of 9.243x10^-4^±3.322x10^-4^ μm^3^ in a 95% confidence interval between [9.022x10^-4^; 9.022x10^-4^] μm^3^ while tubular PVs average a volume of 1.067x10-3±3,322x10^-4^ μm^3^ with a 95% confidence interval between [9.843x10^-4^; 1.150x10^-3^] μm^3^, a statistically significant difference (t-student test, (p< 0.0001), corroborating PV grouping in these two classes. To further investigate morphological heterogeneity of PVs, we analysed trophozoite ultrastructure using freeze fracture scanning electron microscopy. We documented PV heterogeneity and the presence of spherical and tubular PV forms (Supplementary Figure 2). Additional ultrastructural studies using transmission electron microscopy (TEM) were consistent with this classification (Supplementary figure 3A and B).

We proceeded with the rendering of 269 SVs – small spherical vesicles, with distinctly higher electron density than PVs and what could be a coat on the cytoplasmic side of the delimiting membrane (figure 1G). SVs were also identified by TEM (Supplementary figure 3), proximal to the PM. SVs average a volume of 2.525x10^-4^±9.280x10^-5^ μm^3^ in a 95% confidence interval between [2.414x10^-4^; 2.637x10^-4^] μm^3^ (figure 1G, box-plot on the left). This equals to an average diameter of 77.23±9.666 nm in a 95% confidence interval between [76.07; 78.40] nm, differing significantly from spherical PVs which average 120.1±9.507 nm in a 95% confidence interval between [119.2; 121] nm (p< 0.0001) (figure 1G, box-plot on the right). Thus, there is statistical support for SVs as a distinct category of membrane-bounded vesicles (Supplementary figure 3A and C).

Taken together, these findings lead us to hypothesize that, unlike previously thought, PVs are morphologically heterogenous and may comprise different functional categories (Poteryaev et al. 2010; Hipolito et al. 2018; Suresh et al. 2020). However, these data are currently insufficient to determine whether distinct morphologies correlate with distinct functions.

### Combining super-resolution microscopy with ML-assisted image analysis identifies three classes of endocytic compartments in *G. lamblia* trophozoites

FIB-SEM as a technique is not well-suited to the investigation of large cell numbers, and TEM cannot readily provide 3D volumetric information on subcellular compartments. Hence, to address PV heterogeneity in more detail, we continued our investigation of *Giardia* endocytic compartments by Super-resolution Light Microscopy (SRM) techniques and ML assisted image analysis of compartment shapes.

The dimensions of *Giardia* endocytic compartments are well below the diffraction limit of conventional light microscopy (Combs and Shroff 2017). To overcome the Abbe diffraction barrier we used stimulated emission depletion microscopy (STED), potentially achieving a lateral (x,y) and axial (z) resolution of 25-50 nm and 60-100 nm, respectively. This technique decreases the point spread function signal from the illuminated region (Klar et al. 2000; Willig et al. 2006) and allows for accurate imaging of trophozoite PV lumina loaded with a highly photostable fluid phase marker (10 kDa Dextran-Alexa Fluor 594) which is readily taken up into PVs via the fluid phase endocytic pathway (Figure 2A and Supplemental Video 2). In addition to spherical and tubular PVs documented in FIB-SEM, using STED we also determined the presence of polymorphic dextran-labelled organelles i.e., spherical PVs with elongated rods (figure 2A). All labelled PVs were further analysed using the ML-assisted algorithm of the Ilastik program suite. We first performed supervised pixel segmentation followed by supervised object classification. In this second step, we defined and trained the classifier in three organelle morphologies: spherical, tubular, and polymorphic. The latter comprised characteristics of both vesicular and tubular classes, generally with spherical centres with attached tubular protrusions (Figure 2B). After organelle classification we measured their projected areas. Spherical organelles (N =1684) have an average projected area of 0.0205±0.0169 μm^2^ with a 95% confidence interval between [0.0197;0.0213] μm^2^. Tubular endocytic organelles (N=835) present an average projected area of 0.0453±0.0278 μm^2^ with a 95% confidence interval between [0.0435;0.0472] μm^2^. Polymorphic organelles (N=400) have an average projected area of 0.0981±0.0429 μm^2^ with a 95% confidence interval between [0.0939;0.102] μm^2^. ANOVA analysis reveals that each of the three categories is indeed significantly distinct (p<0.0001) based on projected surface area (Figure 2C). This lends further support to the possibility that PV morphological heterogeneity may have functional implications.

**Figure 2.**
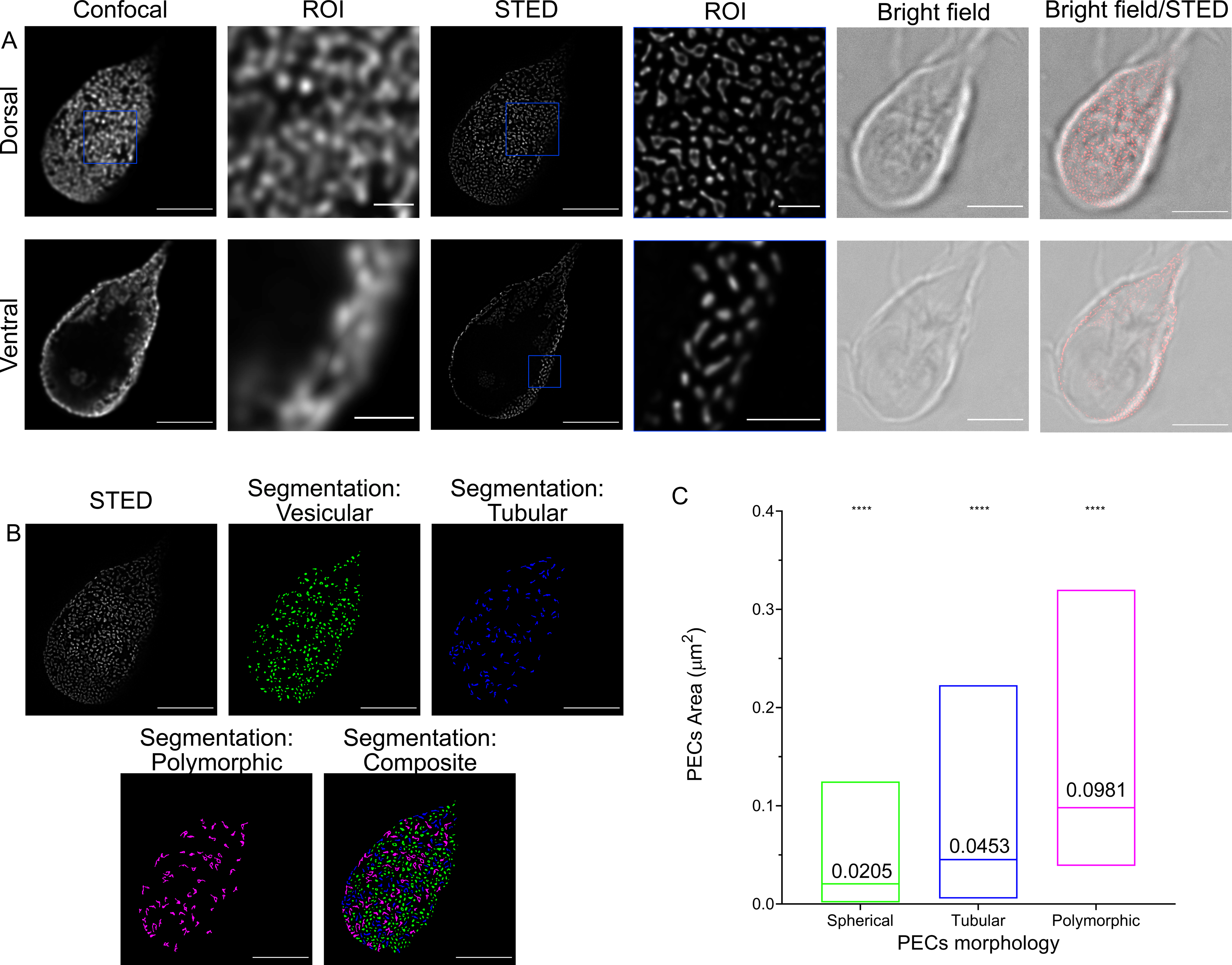
Super Resolution imaging of *Giardia lamblia* peripheral vesicles with Stimulated Emission Depletion (STED). (A) *Giardia* trophozoites loaded with 10kDa Dextran-AlexaFluor 594 were imaged using Confocal and STED microscopy. Dorsal (upper row) and ventral (lower row) regions are represented. In contrast to confocal imaging, STED microscopy allows to separate individual organelles and to visualise different endocytic compartment morphologies. ROI: region of interest. (B) Organelles segmentation with ilastik distinguishes three Dextran-labelled PV categories. (C) PV areas were calculated post-segmentation on maximum projections of the dorsal regions of ≥10 cells, using ilastik. Spherical PVs (green, N =1684) have an average area of 0.0205±0.0169 μm^2^ in a 95% confidence interval between [0.0197;0.0213], ubular PVs (blue, N=835) have an average area of 0.0453±0.0278 μm^2^ in a 95% confidence interval between [0.0435;0.0472]. and polymorphic PVs (magenta, N=400) have an average area of 0.0981±0.0429 μm^2^ in a 95% confidence interval between [0.0939;0.102]. The differences in area are statistically significant (ANOVA; p-value <0.0001). Scale bars: (A) 5 μm and 1 μm (ROIs), (B) 5 μm.

**Figure 3.**
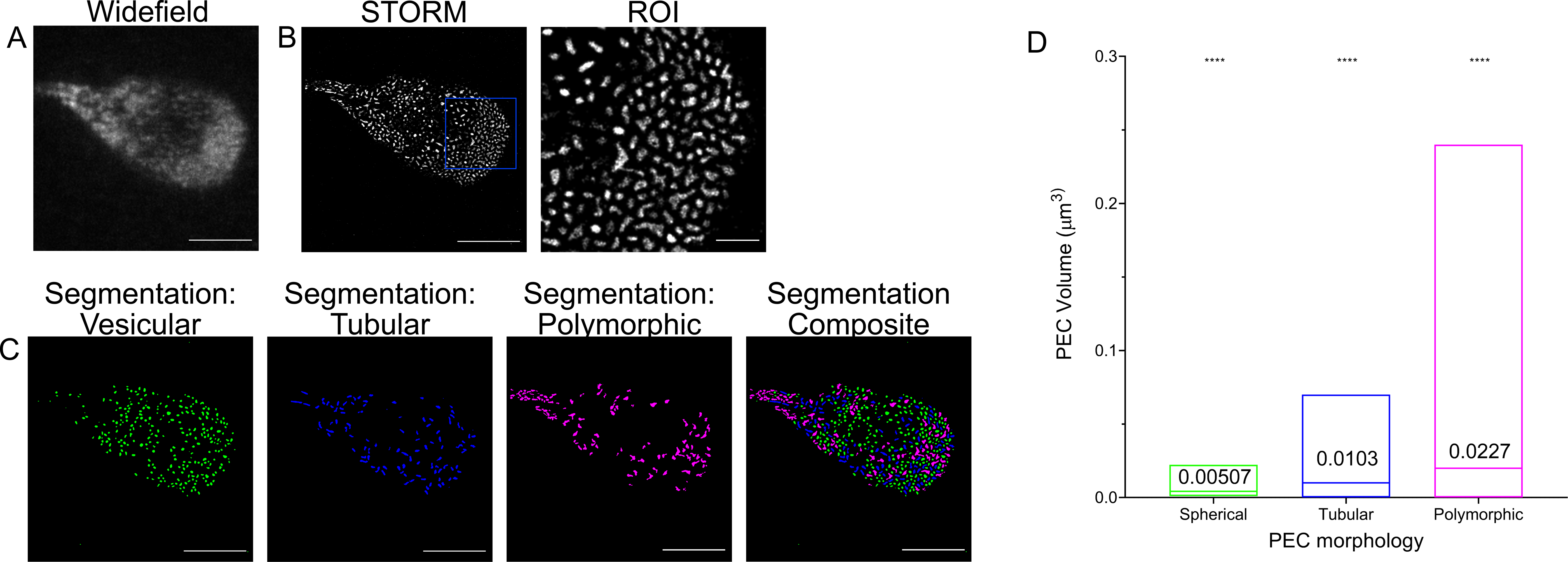
Super Resolution imaging of *Giardia lamblia* PVs by Stochastic Object Reconstruction Microscopy (STORM). (A) Widefield microscopy-based imaging of a Giardia trophozoite loaded with 10kDa Dextran-Alexa Fluor 647. (B) Reconstruction of single molecule events using the Fiji plugin Thunderstorm. As with STED imaging, different PV morphologies are observed. (C) PVs were segmented with the help of ilastik and volumes were calculated and plotted in (D). Spherical PVs (green, N=1989) present an average volume of 0.00507±0.00336 μm^3^ with a 95% confidence interval between [0.00492;0.00522] μm^3^, tubular PVs (blue, N=838) present an average volume of 0.0103±0.00925 μm^3^ in a 95% confidence interval between [0.00967, 0.0109] μm^3^. Polymorphic PVs (magenta, N=1494) present an average volume of 0.0227±0.0214 μm^3^ in a 95% confidence interval between [0.0216, 0.0238] μm^3^. The differences in area are statistically significant (ANOVA; p-value <0.0001). Based on these and previously shown data, the renaming of PVs to Peripheral Endocytic Compartments (PECs), is proposed. Scale bars: 5 μm and 1 μm (ROI),.

Although a STED microscopy-based approach clearly allows resolution of individual organelles as small as PVs, the distinctly lower axial resolution remains limiting for three-dimensional rendering of organelles. Therefore, to push the boundaries of resolution and to further characterize PV morphology, we employed Single Molecule Localization Microscopy (SMLM) (Kao et al. 1994; Huang et al. 2008; Jones et al. 2011).

*Giardia* PVs in trophozoites were loaded with a 10 kDa Dextran-Alexa Fluor 647 fluid phase marker with a high degree of photostability to survive repeated cycles of photoactivation and excitation in SMLM experiments (Dempsey et al. 2011; Olivier et al. 2013). After acquisition, images were reconstructed using the ImageJ plugin ThunderStorm which performs signal centroid calculation, image reconstruction and output (Schindelin et al. 2012; Ovesný et al. 2014). Dextran uptake in PVs was confirmed using conventional widefield microscopy (Figure 3A). STORM image reconstruction shows the subcellular distribution of the fluorescent marker and defines individual organelle lumina (Figure 3B). A closer inspection revealed the presence of morphologically distinct endocytic organelles as previously observed in our FIB-SEM and STED datasets (Figure 3B, ROI and Supplementary Video 3). We again used the supervised ML-assisted algorithm in Ilastik to classify the different morphologies. After a pixel segmentation routine, we performed object classification using supervised ground truth training on subsets of organelles images. Three categories of PVs were defined: spherical, tubular and polymorphic (Figure 3C). To test whether the morphological categorization was consistent with categorization based on organelle volume, we calculated the average lumina volumes of >4000 organelles from the three PV categories. ANOVA testing of organelle volumes for vesicular (0.00507±0.00336 μm^3^, N=1989, 95% confidence interval: [0.00492;0.00522] μm^3)^, tubular (0.0103±0.00925 μm^3^ N=838, 95% confidence interval: [0.00967, 0.0109] μm^3^), and polymorphic organelles (0.0227±0.0214 μm^3^ N=1494, 95% confidence interval: [0.0216, 0.0238] μm^3^ confirmed statistically significant (p <0.0001) morphological differences (Figure 3D and summarised in supplementary table 1).

Taken together, the data generated using three distinct imaging techniques clearly demonstrates PV heterogeneity which may be linked to distinct functions and/or maturation states in this unique endocytic system. To reflect this novel finding and considering that these endocytic and peripherally localized organelles are neither proper vesicles nor canonical vacuoles, we propose renaming PVs to peripheral endocytic compartments (PECs).

### Comparative analysis of endocytic and secretory organelles in *Giardia*, *Spironucleus* sp. and *T. foetus*

*Giardia* spp. have evolved a unique cell architecture including a dedicated organelle for attachment to the small intestinal epithelium – the ventral disk (VD) (Dawson 2010; Brown et al. 2016). In turn, this innovation defines a distinct dorso-ventral as well as antero-posterior polarization of the flagellated trophozoite, marked by swimming directionality. PVs/PECs localize exclusively to the dome-shaped dorsal parasite PM except a small circular patch at the centre of the VD called the bare zone (Zumthor et al. 2016; Cernikova et al. 2020). The result is a maximally decentralized architecture of the *Giardia* endocytic system forming a single-layer interface of what we now appreciate as 3 morphologically distinct organelle classes between the cell exterior and the cytoplasm/ER (Abodeely et al. 2009; Zumthor et al. 2016). We asked whether this type of decentralized sub-PM localisation and polymorphic morphology of endocytic compartments was also represented in other tractable related members of the Diplomonadida and phylogenetically more distant metamonada lineages which do not have a VD, respectively *Spironucleus vortens* and *Spironucleus salmonicida* and the parabasalid *Tritrichomonas foetus*

The diplomonads *Spironucleus vortens* and *Spironucleus salmonicida* are amongst the closest tractable relatives of *G. lamblia* that can be grown axenically under similar conditions (Paull and Matthews 2001; Jørgensen and Sterud 2007; Kolisko et al. 2008a; Xu et al. 2014; Xu et al. 2016). Their endocytic compartments and machineries are partially characterized, with some reports of large vacuolar structures detected by electron microscopy in trophozoites (Sterud and Poynton 2002; Ástvaldsson et al. 2019). Unlike *Giardia*, both species lack dorso-ventral polarization but display a distinct antero-posterior axis. Putative endocytic organelles in *S. vortens* have been detected by fluorescence microscopy of live and fixed cells after incubation with fluorophore-coupled dextran (Zumthor et al. 2016). To further investigate these endocytic compartments, we incubated *S. vortens* and *S. salmonicida* trophozoites with a 10 kDa Dextran-TexasRed fluid phase marker (Figure 4). In stark contrast to the distinctly arrayed PV/PEC labelling seen in *Giardia lamblia* (Figure 4A), confocal microscopy revealed the presence of several dispersed labelled organelles in both *S. vortens* (Figure 4B) and *S. salmonicida* (Figure 4C). *Spironucleus spp.* display several relatively large globular membrane compartments, similar to those observed in well-characterized model organisms (Huotari and Helenius 2011; Day et al. 2018) lacking a fixed subcellular localization. While *S. salmonicida* endocytic compartments localise mostly at the cell periphery (Figure 4C), *S. vortens* organelles present both peripheral and central localizations (Figure 4B and Supplementary Video 4). We also assessed endosome morphology in *T. foetus* using the same labelled dextran-based approach. Similar to *Spironucleus* species, *T. foetus* presents an antero-posterior axis but no attachment organelle nor dorso-ventral polarization. Similar to *Spironucleus spp.*, *T. foetus* accumulated the endocytosed fluid phase maker in several globular endocytic compartments (Figure 4D) consistent with previous reports on vacuolar structures identified in *T. foetus* by electron microscopy (Lealda et al. 1986). In our ultrastructure observations we could not detect multivesicular bodies or vacuoles containing intra-luminal vesicles in either *Spironucleus* spp. or *T. foetus*.

**Figure 4.**
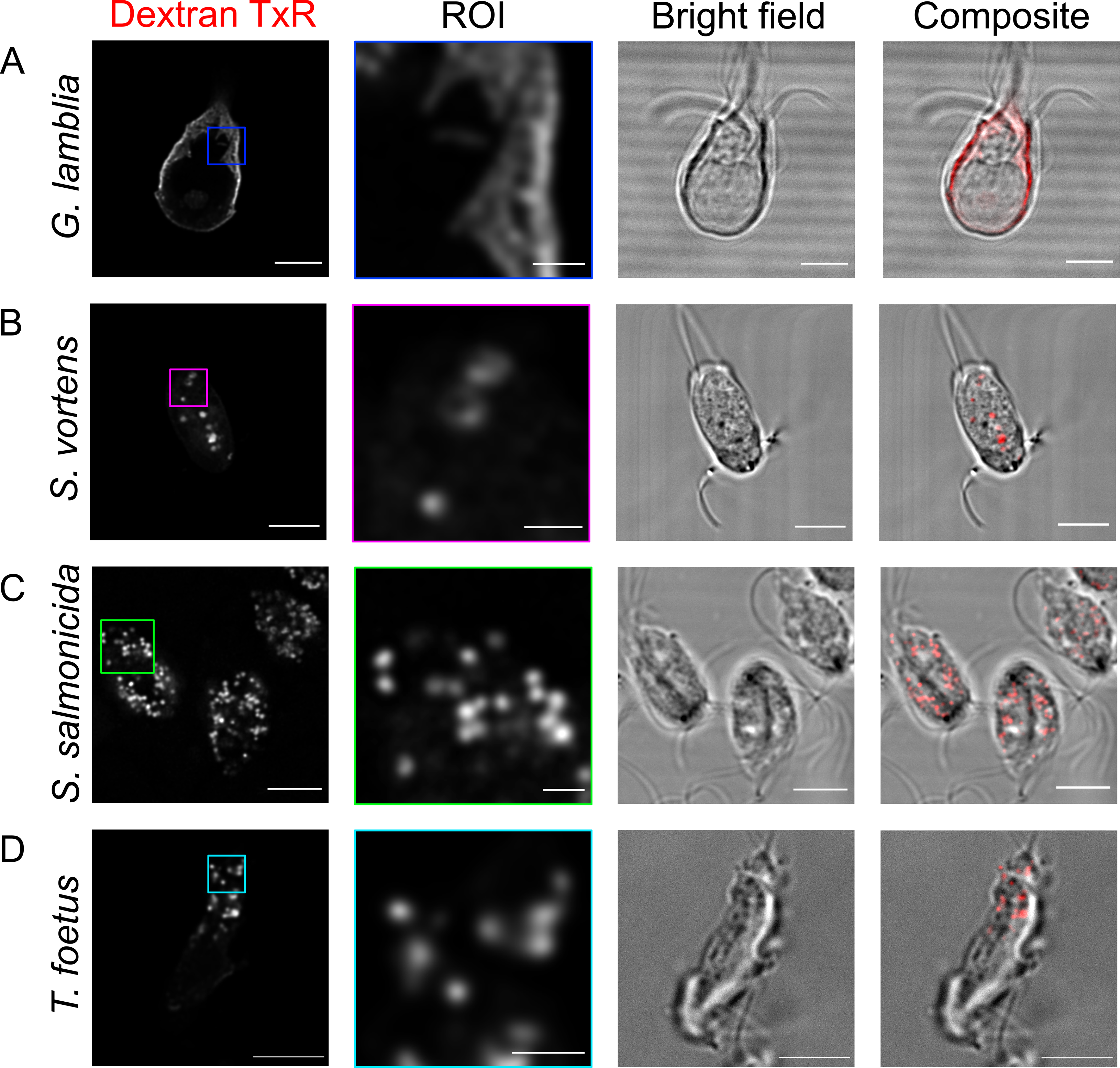
Uptake of fluorescently-labelled dextran in metamonada and discoba members: *G. lamblia*, *S. vortens*, *S. salmonicida* and *T. foetus* after 30 minutes. (A) *G. lamblia* cells present endocytic compartments spread on the cell periphery not resolved by conventional light microscopy. On the other hand, (B) *S. vortens*, (C) *S. salmonicida* and (D) *T. foetus* present vesicular shaped endocytic compartments. Scale bars: 5 μm (full cells) and 1 μm (ROI).

Taken together, these data show how, in closely-related protozoa lacking dorso-ventral polarization and a dedicated attachment organelle, endocytic organelles appear to have no specific localization. This lends support to the notion that PV organelle architecture is intimately associated to the emergence of the VD, both as adaptations to the mammalian small intestine niche (Zumthor et al., 2016).

To visualize and measure morphological parameters of *Spironucleus* and *T. foetus* endocytic compartments, we performed 2D-STED imaging and transmission electron microscopy (TEM). *S. vortens* cells loaded with 10 kDa Dextran-Alexa Fluor 594 showed accumulation of the fluid phase marker in roughly spherical organelles (Figure 5A). Labelled endocytic vacuoles have an average diameter of 468±206 nm (95% confidence interval [421;515] nm, N=10) (Figure 5A, box-plot). Volumetric rendering of 3D reconstructed optical sections document the uniformly globular morphology of these organelles (Supplementary Video 5). TEM analysis revealed an ellipsoid shape of endocytic organelles in *S. vortens* with an average maximal diameter of 844±335 nm in a 95% confidence interval between [763;905] nm (Figure 5B, box-plot). The dimensions measured in TEM represent those of the membrane-delimited organelle. In contrast, the dimensions measured by STED represent a projection of the fluid phase marker distribution within the available organelle lumen. The fact that the former (844±335 nm) is larger than the latter (468±206 nm) indicates that these organelles contain additional cargo which prevents the endocytosed fluid phase marker to distribute in the complete compartment volume delimited by the organelle membranes. TEM investigation in *S. salmonicida* cells (Supplemental Figure 4A) showed the presence of small globular vacuoles (V) close to the PM (Supplemental Figure 4B) with an average diameter of 205±62.6 nm (N=114) in a 95% confidence interval of [193;217] nm (Supplemental Figure 4E). These vacuoles are smaller than the ones found in *S. vortens* (Supplementary Figure 4C,D,F, p-value < 0.0001). In these conditions, neither coated vesicles nor a stacked Golgi apparatus could be documented in *S. vortens* or *S. salmonicida*.

**Figure 5.**
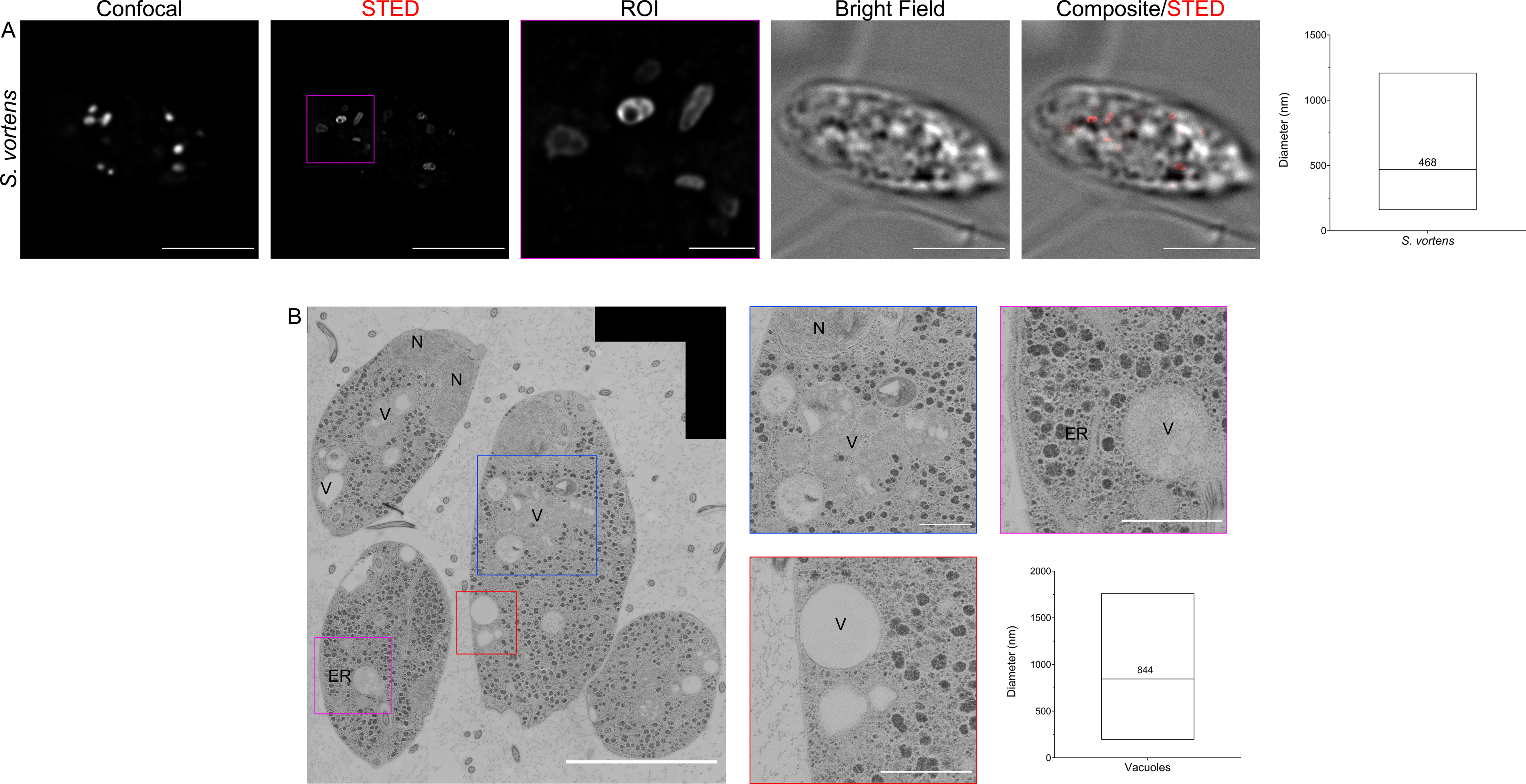
Super Resolution imaging of *S. vortens* endocytic compartments with STED and Transmission Electron Microscopy (TEM) (A) Following incubation with dextran-TexasRed for 30 minutes, *S. vortens* display elongated endocytic compartments with an average diameter of 468±206 nm in a 95% confidence interval between [421;515] nm (upper box-plot). (B) TEM imaging detects several endosome-like vacuoles (V) throughout the cell, with an average diameter of 844±335 nm in a 95% confidence interval between [763;905] nm (lower box-plot). Measurements were done manually. The Endoplasmic Reticulum (ER) and a nucleus (N) are also highlighted in the images. Scale bars: 5 μm (full field of view) and 1 μm (ROIs).

2D-STED analysis of *T. foetus* cells incubated with 10 kDa Dextran-Alexa Fluor 594 revealed a roughly circular distribution of the marker within endocytic vacuoles (Figure 6A and Supplemental Video 6) with an average maximal diameter of 517±251 nm (95% confidence interval [455;580] nm, N=10) (Figure 6A, box-plot). TEM imaging revealed the presence of two distinct classes of endosome-like vesicles (Figure 6B) based on electron density of the lumen. Low-density vesicles were identified both at the cell periphery and in central areas termed vacuoles (V); vesicles of higher electron density were previously identified as digestive vacuoles (DVs) (Lealda et al. 1986) and contain structured material and membranes. Analysis of TEM micrographs showed that DVs are significantly larger than vacuoles, with an average diameter of 764±203 nm (N=50) (95% confidence interval [707;822] nm). Vacuoles in turn have an average diameter of 246±100 nm (N=153) in (95% confidence interval [230;262] nm) (Figure 6B, box-plot). Stacked Golgi organelles are abundant in TEM micrographs of *T. foetus* trophozoites, as documented previously (Rosa et al. 2014) (Supplemental Figure 5A). Consistent with a more canonical architecture of the membrane trafficking system in *T. foetus,* coated vesicles were observed in the cytosol particularly in the vicinity of Golgi stacks (Supplemental Figure 5B) (Lealda et al. 1986; Midlej et al. 2011; Schlacht et al. 2014). These vesicles averaged a diameter of 58.4±13.1 nm (N=128) (95% confidence interval [56.1;60.7] nm) corresponding to the size range of clathrin coated vesicles (CCVs) (Traub 2011).

**Figure 6.**
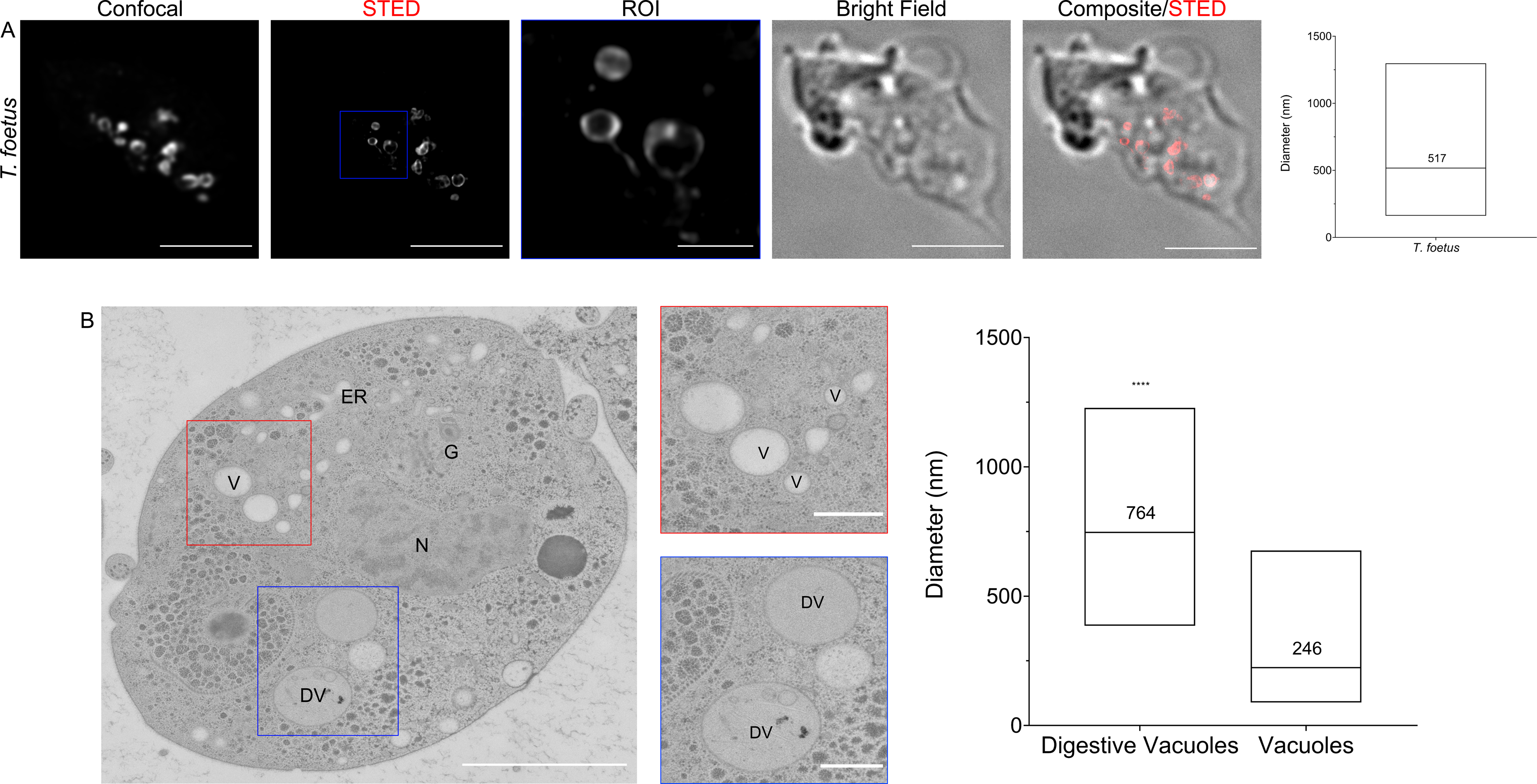
Super Resolution Imaging of *T. foetus* endocytic compartments with STED and TEM. (A) 2D-STED analysis of *T. foetus* cells loaded with 10 kDa Dextran-Alexa Fluor 594 illuminate endosomes with globular structures at an average diameter of 517±251 nm in a 95% confidence interval between [455;580] nm (upper box-plot). (B) TEM investigation of *T. foetus* cells reveals two kinds of endosome-like vesicles. Low electron-density vesicles found both at the cell periphery and centre are deemed vacuoles (V) with an average diameter of 246±100 nm (N=153) in a 95% confidence interval of [230;262] nm. Vesicles of higher electron-density and with noticeable content are Digestive Vacuoles (DVs). DVs are larger than vacuoles, with an average diameter of 764±203 nm (N=50) in a 95% confidence interval of [707;822] nm (lower box-plot). Scale bars: 5 μm (full field of view) and 1 μm (ROIs).

Finally, to probe the dynamics of endocytic compartments in *G. lamblia*, *S. vortens*, *S. salmonicida* and *T. foetus*, cells were exposed to 10 kDa Dextran-TexasRed for 5, 10, 20 or 30 minutes, fixed chemically, and imaged by confocal microscopy (Figure 7). The number of *G. lamblia* PECs labelled with the fluid-phase marker increased over time, with the label accumulating strictly at the cell periphery (Figure 7A). In contrast, endocytic compartments in *S. vortens* were first visualized at the PM, and were then observed at more central locations of the cell at later time points. Given the overall increase in fluorescent intensity and the motile nature of these organelles, it appears there is a constant uptake of dextran over the analysed period (Figure 7B). In these conditions, *S. salmonicida* and T. foetus vacuoles behave similarly since both sets of organelles are diffused within the cell cytoplasm, with a steady decrease and then marked increase in dextran content (Figure 7C and D).

**Figure 7.**
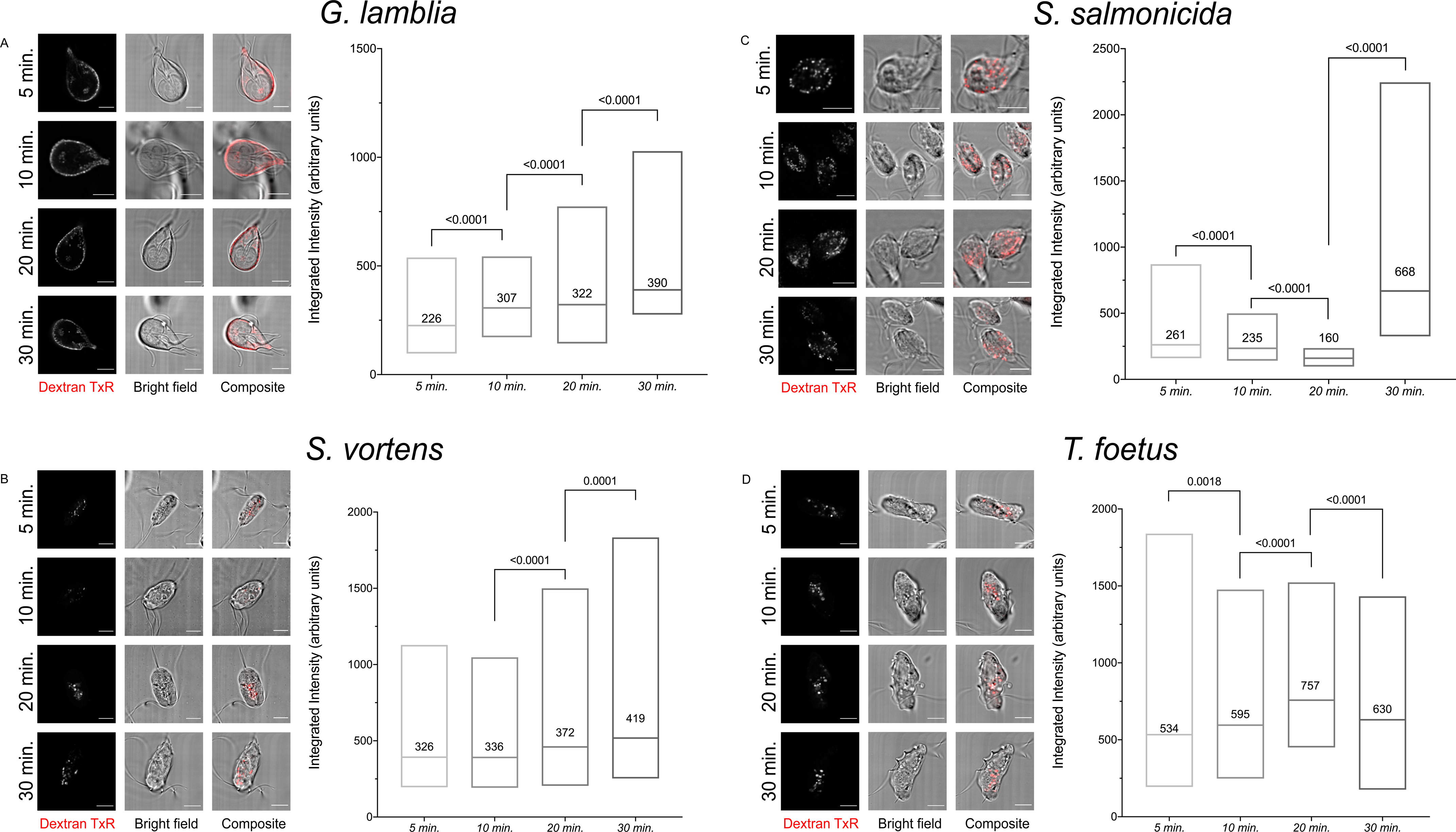
Time course on fluorescent dextran uptake in selected metamonada and discoba specimens: *G. lamblia*, *S. vortens*, *S. salmonicida* and *T. foetus* after 30 minutes. **(A)** PV/PECs in *G. lamblia* start acquiring external material right after 5 minutes of incubation with dextran. Over 30 minutes of dextran incubation, the endocytic marker does not leave PECs. In (B) *S. vortens*, (C) *S. salmonicida* and (D) *T. foetus,* dextran is up taken in small vesicles. In *S. vortens* these small vesicles tend to agglomerate at the centre of the cell while in *S. salmonicida*, these vesicles seem to stay peripheral. In *T. foetus*, vesicles bearing dextran from at the periphery of the cell and also migrate to the interior of the cell. All scale bars: 5 μm.

### *Gl*CHC foci associate with different classes of giardial PECs with different stoichiometry

Previously, we established that *Giardia* clathrin heavy chain (*Gl*CHC) associates to discrete static foci at the dorsal PM of trophozoites, in close proximity to PV/PECs. Further, *Gl*CHC strongly interacts with a putative albeit highly diverged *Giardia* clathrin light chain homologue previously named *Gl*CLC (Zumthor et al. 2016; Cernikova et al. 2020). We wondered whether association to clathrin assemblies holds any relation to the heterogeneity of PV morphology. To address this question, we used STED microscopy to investigate epitope-tagged *Gl*CHC (*Gl*CHC-HA) deposition at distinct foci at the dorsal PM and the cell periphery, consistent with PV/PECs location (Figure 8A). Segmentation of foci using a ML-assisted Ilastik tool, allowed to determine the dimensions of *Gl*CHC at an average diameter of 134±36.6 nm (N = 4524) (95% confidence interval [132;135] nm) (Figure 8B). Similar to GlCHC, subcellular distribution of epitope-tagged *Gl*CLC (*Gl*CLC-HA) showed an identical pattern consistent with its demonstrated direct interaction with *Gl*CHC (figure 8C) (Zumthor et al. 2016). Segmentation of foci using a ML-assisted Ilastik tool, determined the dimensions of *Gl*CLC foci at an average diameter of 159±48.8 nm (N = 984) (95% confidence interval [156;162] nm) (Figure 8D), Notably, the average size of *Gl*CLC foci is larger than that of *Gl*CHC foci (p< 0.0001, t-student test) (Figure 8E).

**Figure 8.**
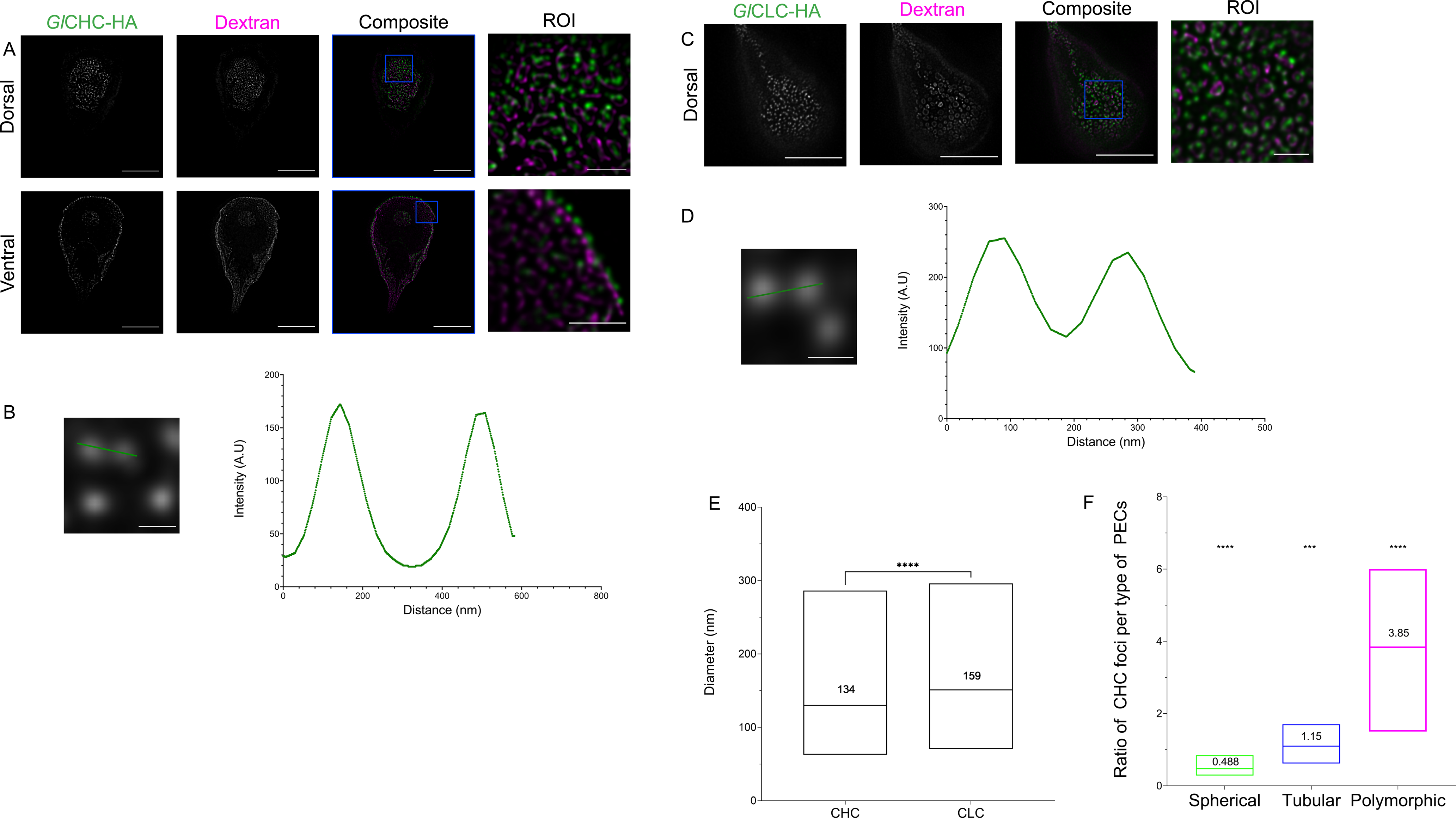
*Gl*CHC and *Gl*CLC foci associate to *Giardia* PECs with different stoichiometries. (A) An epitope-tagged *Gl*CHC reporter was used to localise foci of GlCHC deposition at the cell periphery beneath the PM, by STED microscopy.(B) These foci can be clearly resolved with STED microscopy (left graph). (C) The same was done with an epitope-tagged *Gl*CLC reporter and (D) foci resolved as in (B) (right graph). (E) Foci were segmented with ilastik. Areas were determined and diameters calculated in an automatic procedure assuming spherical geometry. *Gl*CHC foci present an average diameter of 134±36.6 nm (N = 4524) in a 95% confidence interval between [132;135] nm. *Gl*CLC foci average a diameter of 159±48.8 nm (N = 984) in a 95% confidence interval between [156;162] nm. *Gl*CLC foci are larger than *Gl*CHC in significant statistical manner (p-value < 0.0001, t-student test). (F) *Gl*CHC foci associate in different stoichiometry to different classes of PECs. Spherical PECs are either not associated to clathrin or with just one focus: mean of 0.488±0.159 foci per spherical in a 95% confidence interval between [0.403;0.573] foci per spherical organelle. Tubular PECs associate with one focus of *Gl*CHC: mean of 1.15±0.287 in a 95% confidence interval between [0.994;1.3] foci per tubular organelle. Polymorphic PECs associate with 3 or more *Gl*CHC foci with an average of 3.85±1.14 in a 95% confidence interval between [3.25;4.46] foci per polymorphic organelle. These distributions are statistically significative with a p-value < 0.0001 (ANOVA analysis).

Using STED microscopy, we further determined the number of *Gl*CHC foci showing signal overlap with the 3 classes of Dextran-Texas Red loaded PV/PECs (figure 8F). By calculating the degree of signal overlap between *Gl*CHC foci and PEC lumina, we determined that spherical PECs are associated with at most one *Gl*CHC focus with an average of 0.488±0.159 foci per spherical PEC (95% confidence interval [0.403;0.573]). Tubular PECs associated with at least one *Gl*CHC focus with an average of 1.15±0.287 foci per tubular PEC (95% confidence interval [0.994;1.3]). Polymorphic PECs associated with 3 or more *Gl*CHC foci with an average of 3.85±1.14 foci per PEC (95% confidence interval [3.25;4.46]). Taken together, we find a directly proportional and statistically-significant ratio of clathrin foci to PEC size and type (ANOVA; p-value < 0.0001). This is in line with the notion that PV morphological heterogeneity is indeed correlated with organelle functional diversity, as measured by association to clathrin assemblies.

### Pan-eukaryotic searches for CHC and CLC reveal loss of a *bona fide* CLC within the fornicata lineage and the emergence of putative CLC analogues

We previously proposed that PV/PEC localization at the dorsal PM of trophozoites and evolution of the adhesive disc attachment organelle are interdependent adaptations to life in direct contact with the host’s gut epithelium (Zumthor et al. 2016). Furthermore, we found a direct correlation between types of PV/PEC and the number of foci of clathrin assemblies. Given that the role for clathrin assemblies in Giardia has not been elucidated and that the nature of *Gl*CLC (*Gl*4259) as a clathrin light chain orthologue is dubious, we sought to shed light on the significance of this correlation by investigating the distribution of both CHC and CLC orthologues in selected eukaryotic lineages. To do this, we employed protein homology searches based on Hidden-Markov Models (HMM) (Eddy 2011) using as query an alignment of canonical and documented CHC or CLC sequences from several protozoa and metazoan species (Supplementary Tables 2 and 5) (Morgan et al. 2001; Kaksonen et al. 2005; Adung’a et al. 2013; Kirchhausen et al. 2014; Karnkowska et al. 2016; Karnkowska et al. 2019, Füssy et al., 2021). In our search we considered assembled read data from RNA-seq experiments (transcriptomics) as reliable as genomic sequence data (Cheon et al. 2020). In this case, we used the reference CHC or CLC sequences and performed tblastn searches. Nucleotide sequences from each reliable hit (lowest e-value) were translated and subjected to a reciprocal blast-p analysis to validate protein identity. We found CHC homologues in all selected genomes and transcriptomes we searched, highlighting the likely essential nature of CHC (Figure 9A and supplementary table 4 and 5).

**Figure 9.**
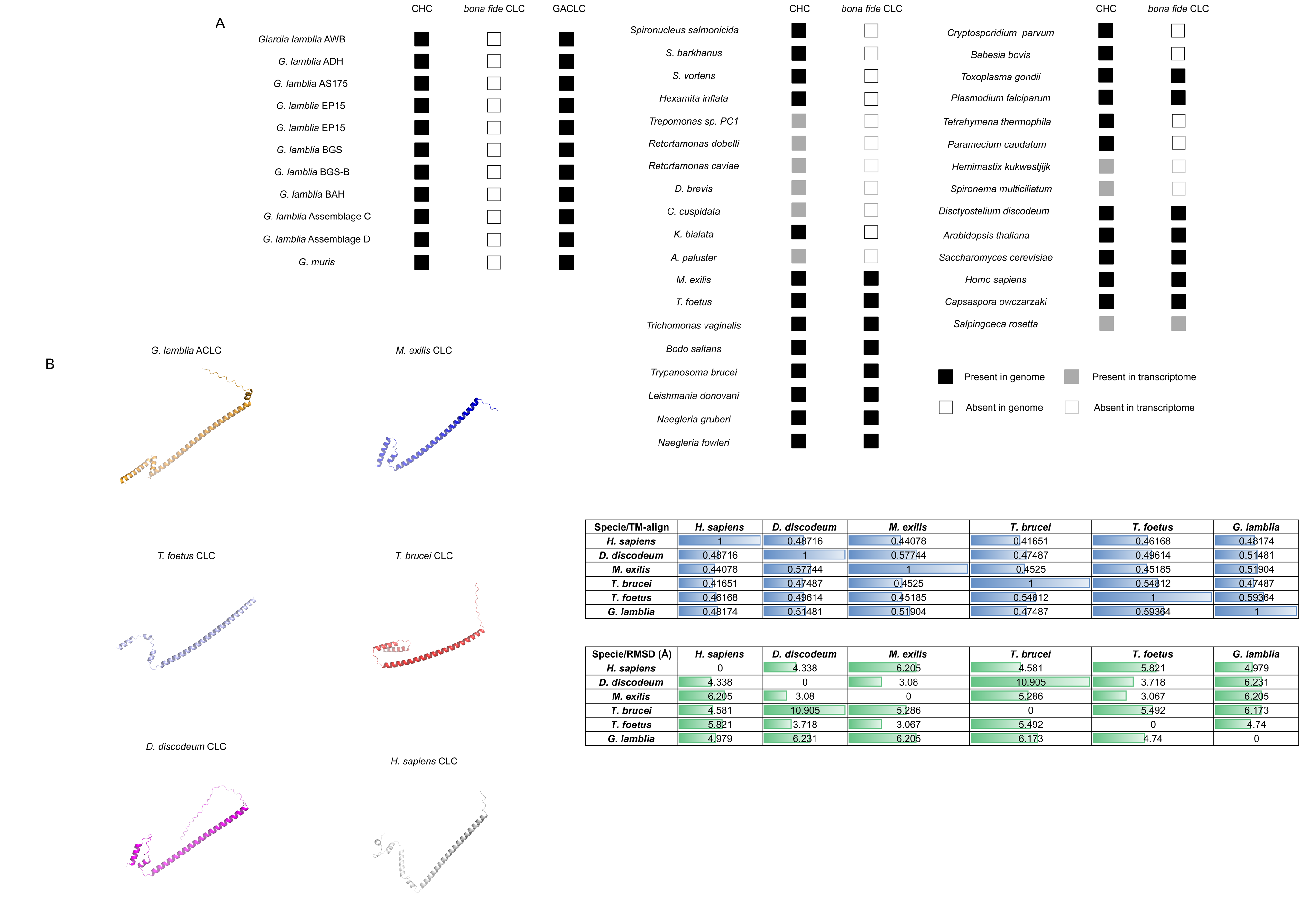
Homology search for *bona fide* CHC and CLC reveal key importance of CHC and patchy conservation of CLC. (A) CHC and CLC homology searches demonstrate the conservation of CHC orthologs in all analysed lineages. However, several lineages appeared to have lost a readily-detectable *bona fide* CLC including all selected Fornicata, certain Chromerids (Woo et al. 2015), some ciliates (Richardson and Dacks 2022) and members of the Hemimastigophora. (B) *Ab initio in silico* protein modelling of *G. lamblia* ACLC (formerly *Gl*CLC/Gl4529), *T. brucei* CLC, *T. foetus* CLC, *M. exilis* CLC, *D. discoideum* and *H. sapiens* CLC sequences using AlphaFold, the current standard in *in silico protein* structure modelling. TM-align and RMSD values were calculated showing close structure analogy between predicted structures (table).

*Gl*CHC is a clearly divergent ortholog compared to its counterpart in eukaryotic model organisms, with only 24% amino acid identity to human CHC (Marti, Regös, et al. 2003). A domain analysis of selected CHC sequences (supplementary figure 6 and supplemental table 10) reveals that *Gl*CHC contains fewer α-helical domains than other analysed CHC sequences, further highlighting its divergence. We also performed an in-depth search for the CHC triskelion uncoating “QLMLT” motif which we documented previously to be missing in *Giardia* (Fotin et al. 2004; Rapoport et al. 2008; Zumthor et al. 2016). Notably, this motif appears to be only present in Metazoa and in the closely related Filastera and Choanoflagellata (King et al. 2008; Fairclough et al. 2013; Suga et al. 2013). In Fungi only a partial “L(M)TL”motif was identified and we were unable to detect a conserved uncoating motif in CHC sequences of members of the Archaeplastida, Amoebozoa or SAR supergroups (supplementary figure 7).

In stark contrast to CHC, the search for *bona fide* CLC sequences did not retrieve any reliable predictions in available genomes and transcriptomes from species of the Fornicata lineage, including the lineages Hexamitidae, Retortamonas and Carpediemonas-like organisms (Xu et al. 2014; Leger et al. 2017; Tanifuji et al. 2018; Füssy et al. 2021; Salas-Leiva et al. 2021). Importantly, this search did not return the putative, highly diverged *Gl*CLC (Zumthor et al. 2016).

There are documented CLC orthologues in members of the Discoba, such as *Trypanosoma brucei* CLC (Tb927.10.14760) (Manna et al. 2017) and in the parabasalid *Trichomonas vaginalis* (TVAG_29749) (Carlton et al. 2007; Aurrecoechea et al. 2009). Furthermore, we readily identified a CLC homologue in *T. foetus* (gene accession OHT14195.1, forward HMMer e-value of 1.00E-26 and reverse Blastp e-value of 2.00E-11, returning the human CLC homologue) (Figure 9A). Therefore, while *bona fide* CLC orthologues can be readily identified in Discoba and Euglenozoa members and Preaxostyla – as in the metamonad *Monocercomonoides exilis* (Karnkowska et al. 2016) – no sequence could be found among members of the Fornicata lineage. Further, we were unable to identify a *bona fide* CLC sequence within the newly documented transcriptome of Hemimastigophora (Lax et al. 2018). These observations are in line with the notion that, unlike CHC, CLC is dispensable. This is also supported by failures to identify *bona fide* CLC in chromerids such as *Cryptosporidium parvum* and *Babesia bovis* (Woo et al. 2015) and in some ciliate lineages, such as *Tetrahymena thermophila* and *Paramecium caudatum* (Richardson and Dacks 2022).

Given that *Gl*CLC’s predicted 3D structure is reminiscent of CLCs (Zumthor et al., 2016) but it could not be retrieved as related to a *bona fide* CLC, with its only known orthologue found in *Giardia muris* (Figure 9A), *Gl*CLC was further analysed using the HHPred suit, in the attempt to find distantly-related non-Giardia sequences (Zimmermann et al. 2017). This search retrieved no robust prediction for a non-Giardia sequence (supplemental table 9). Given that the degree of divergence is such that no reliable claim to orthology can currently be supported and no orthologue for *Gl*CLC can be found outside the *Giardia* genus, we propose the renaming of GlCLC to *Giardia lamblia* Analogous to Clathrin Light Chain-***Gl*ACLC**-as a CLC structural analogue acquired and retained in the last *Giardia* common ancestor (LGCA). This appears to correlate with loss of a *bona fide* CLC with the last Fornicata common ancestor (LFCA).

To test the extent of environmental pressure on this protein family’s evolution, we calculated synonymous vs non-synonymous mutation ratios (ω = ks/kn) for *Gl*ACLC homologues (supplemental figure 8). Interestingly, known sequences for all *Giardia* isolates present a ω < 1 which indicates that current sequences are not under selective pressure to evolve.

To further investigate the structural analogy of *Gl*ACLC to canonical CLCs, we performed *in silico* modelling of its C-terminal domain () using the new standard in *ab initio* protein structure modelling – AlphaFold - based on deep-learning neural networks (Jumper et al. 2021; Tunyasuvunakool et al. 2021) (Figure 9B and supplemental table 11). Template modelling score independent of sequence (Tm-align) and Root Mean Square Deviation (RMSD) (Zhang and Skolnick 2005; Kufareva and Abagyan 2012) values provide substantial evidence for structural analogy of *Gl*ACLC and canonical CLCs, in line with previous observations (Zumthor et al. 2016). The newly predicted structures for *Gl*ACL*C* have a stronger resemblance to the predicted structure of a mammalian clathrin light chain (Wilbur et al. 2010). Altogether, the presented *in silico* data strongly suggest *Gl*ACLC to be structural analogue of CLC.

### *Ss*CHC is distributed in the cytosol and interacts with a putative light chain structural analogue

We hypothesize that *GlA*CLC is a Giardia-specific CLC analogue, which arose *de novo* in the LGCA, possibly to supplant the loss of a *bona fide* CLC, We wondered whether *de novo* acquisition of a CLC analogue with divergent sequence but preservation of structural and potentially also functional features had occurred independently in other Diplomonadida lineages. To address this question, we selected *Spironucleus salmonicida*the closest genetically tractable and sequenced relative to *Giardia* (Jerlström-Hultqvist et al. 2012; Xu et al. 2014) in which a *bona fide* CLC cannot be detected, We expressed an epitope-tagged variant of the *S. salmonicida* CHC orthologue (*Ss*CHC-3xHA) and detect it distributed in a punctate pattern throughout the trophozoite cytosol. Using higher resolution confocal microscopy, *Ss*CHC-3xHA was shown in cytoplasmic structures reminiscent of giardial CHC focal assemblies (Supplemental Figure 9A and B and Supplemental Video 7).

To probe for the presence of a CLC analogue in *Spironucleus*, we performed native co-immunoprecipitation experiments (native co-IP) using the *Ss*CHC-3xHA-expressing *S. salmonicida* transgenic line. *In silico* analysis of the mass spectrometry dataset focused on the most abundant proteins pulled down with a minimum of 10 peptide hits using stringent criteria. This yielded 171 proteins which were either exclusive to the native co-IP sample derived from the transgenic reporter line, or ≥3-fold enriched in the transgenic line compared to the non-transgenic parental strain (Supplemental Table 12). We identified several endocytosis-related proteins, with *Ss*-dynamin being the most abundant (48 hits and exclusive to *Ss*CHC co-IP reaction), together with *Ss*-β-adaptin, *Ss*-calmodulin and *Ss*Sec7 (Supplemental Figure 9C and D). Despite its intranuclear localization, annexin 5 is also found to be a putative interactor of *Ss*CHC (Einarsson et al. 2019).

Since we postulated that a possible CLC analogue would be among the hypothetical proteins, we probed those hits using the HHPred algorithm (Zimmermann et al. 2017), focusing on candidates with predicted secondary structures composed of alpha-helical and coiled-coil domains and a size < 40 kDa, consistent with all CLC documented thus far. One of these, protein *Ss*50377_11905, was found to be prominently pulled down and contains several coil-coil domains, at a predicted weight of 39kDa. The 150 amino acid C-terminus of the protein was modelled in AlphaFold and superimposed with CLC structures (supplementary figure 9E). TM-align values within structure similarity (0.5 or above), and RMSD values of *circa* 5-6 Å make a compelling case for *Ss*50377_11905 to be a *S. salmonicida* CLC analogue, similar to *Gl*CLC. Using *Ss*11905, we also performed homology searches in the available transcriptome of the related diplomonad *Trepomonas sp.* (Kolisko et al. 2008b; Xu et al. 2016). We found a candidate ortholog, TPC1_16039 (forward tblastn e-value of 1E-5 and reverse blastp e-value of 4E-12) (supplementary table 11). Notably, however, searches using *Ss*50377_11905 into the remaining fornicata representatives failed to retrieve candidate homologues, whether *Gl*ACLC or other (data not shown). Protein structure modelling with AlphaFold and super-imposition with *Ss*11905, *Gl*ACLC, *T. brucei* CLC and *H. sapiens* CLC suggest that TPC1_16039 is also a structural CLC analogue, orthologous to *Ss*11905 (supplementary figure 10).

**Figure 10.**
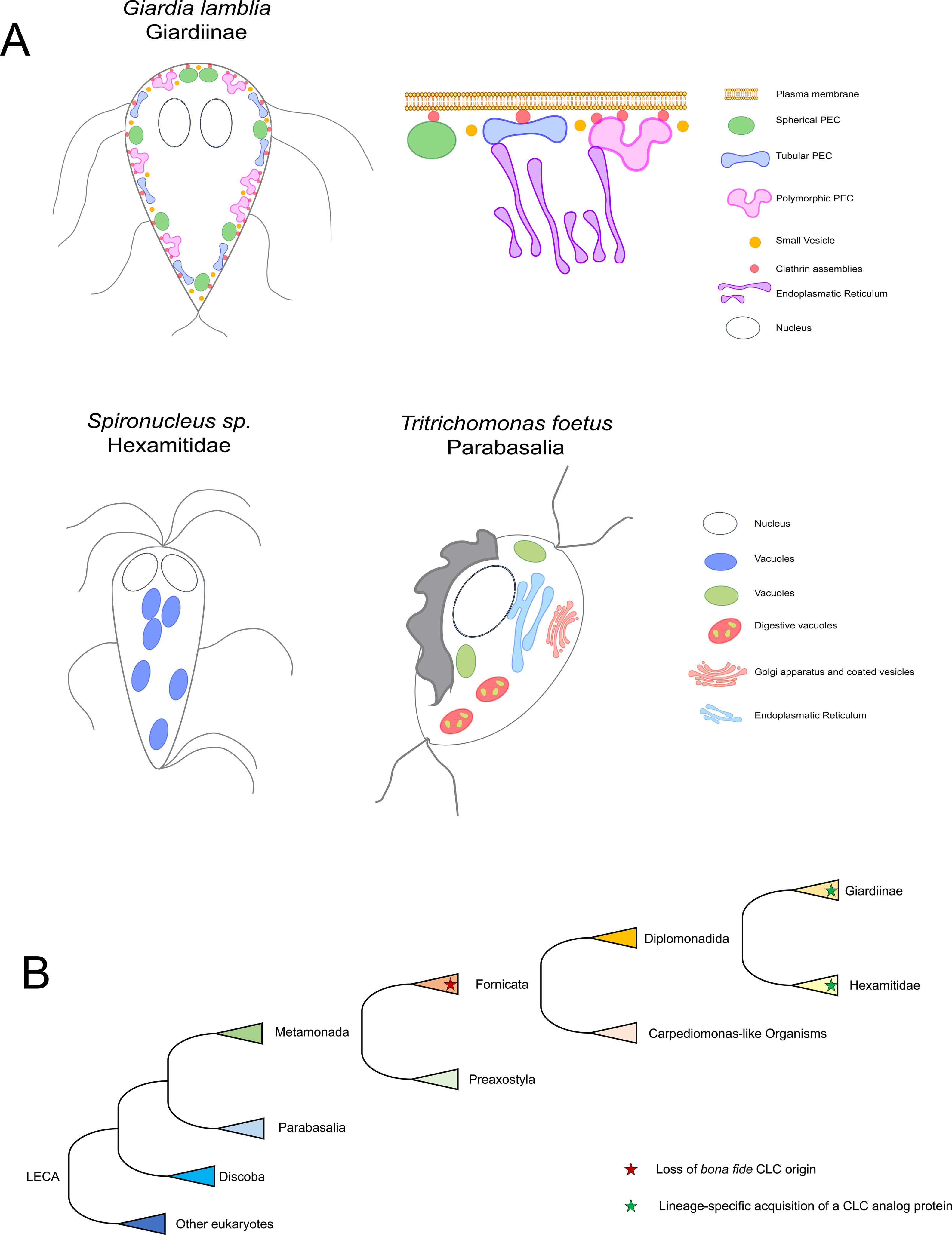
Endosome-like organelle models in *Giardia*, *Spironucleus* and *Tritrichomonas,* and proposed evolution of CLC. (A) Simplified cartoons of the endocytic and secretory pathway in *Giardia lamblia* (Giardiinae), *Spironucleus sp.* (Hexamitidae) and *Tritrichomonas foetus* (Parabasalia). PECs are a hallmark of the Giardia lineage, while more canonical vesicular endosomes are present in both the Spironucleus lineage and Parabasalia. (B) Simplified evolutionary model for *bona fide* CLC. The last eukaryotic common ancestor possessed a *bona fide* CLC which was lost at the last Fornicata common ancestor. In at least two derived lineages, – Giardinae and *Spironucleus* spp. - de *novo protein* analogue to CLC was acquired independently.

## DISCUSSION

### *Giardia lamblia’s* endocytic organelle system consists of three classes of small acidifying membrane compartments

Subsequent to ingestion and excystation, *Giardia* trophozoites attach to the intestinal lumen, proliferating and encysting on localised foci throughout the mucosa of the small intestine (Barash et al. 2017). Nutrients required for this propagation are taken up from the environment through PV/PEC-mediated endocytosis of fluid phase and membrane bound material (Lanfredi-Rangel et al. 1998; Adam 2001; Abodeely et al. 2009; Carranza and Lujan 2009; Cotton et al. 2011; Zumthor et al. 2016; Touz et al. 2018). Despite the essential nature of these endocytic organelles, complete resolution of the ultrastructure of the *Giardia* endocytic pathway remains unsolved. To address this we performed an ultrastructural investigation of *G. lamblia* endocytic compartments to obtain a nanometric view of their morphology as defined by their membrane as well as the lumen accessible to fluid phase markers in labelling experiments (Abodeely et al. 2009; Zumthor et al. 2016).

We began by dissecting an entire *G. lamblia* trophozoite using scanning electron microscopy and focused our analysis on PVs. These structures were segmented and rendered in three dimensions. Using this method unambiguously detected at least two distinct classes of PV morphologies, with some being obviously globular in shape while others presenting a more tubular nature. After expanding our analysis of PVs/PECs to super resolution light microscopy methods STED and STORM (Jacquemet et al. 2020) we determined that PVs/PECs are present in three discernible morphologies: spherical, tubular and polymorphic. Thus, we proposed the renaming of these organelles into peripheral endocytic compartments (PECs).

Furthermore, in line with previous reports (McCaffery et al. 1994; McCaffery and Gillin 1994; Benchimol 2002; Zumthor et al. 2016), we also detected smaller vesicles (SVs) of around 80 nm in radius which appear to be coated, based on their electron-dense surface, and are not related to CHC foci at the PV-PM interface. Identifying the nature of SV coats may shed light on their corresponding cargo. For instance, COPI components such as the small GTPase ARF1 and β’-COPI were found to be located not only at the parasite’s ER but also at the cell periphery (Marti, Regös, et al. 2003; Stefanic et al. 2006; Stefanic et al. 2009), in line with SV distribution. If indeed COPI were found to act as coat for these currently uncharacterized membrane carriers, an intriguing possibility emerges for SVs as vehicles for the trafficking of variant surface proteins (VSPs) to the *Giardia* cell surface. VSP secretion is compromised by the presence of brefeldin-A, implicating ARF-GTPase cycles in VSP trafficking (McCaffery et al. 1994; Lujan et al. 1995; Marti, Li, et al. 2003; Marti, Regös, et al. 2003). Currently, the exact mechanism for VSP translocation from the ER to the plasma membrane remains unknown, although it has been postulated that PVs/PECs may be involved. Furthermore, previous reports exclude the presence of an intermediate VSP trafficking compartment between the ER and the PM detectable by microscopy (Marti, Li, et al. 2003; Marti, Regös, et al. 2003). Given the estimated diameter of SVs at ca. 80 nm, these compartments would have easily escaped detection in standard light microscopy experiments. An alternative hypothesis concerning SVs is that they are peroxisome derivatives. Recently, peroxisomes have been found in *Entamoeba histolytica,* a microaerophile like Giardia, with diameters between 90-100 nm, in the range of *Giardia* SVs (Verner et al. 2021). Furthermore, reports on immuno-EM detection of peroxisome-like proteins in *Giardia* highlighted the presence of small dense vesicles with diameters of *circa.* 100 nm (Acosta-Virgen et al. 2018). Taken together, while we favour the hypothesis of SVs being secretory trafficking vesicles considering their apparent coating, the possibility of SVs corresponding to cryptic peroxisome-like organelles cannot be excluded.

Compared to endosome-like vacuoles in *Carpediemonas*-Like Organisms (CLOs) (Yubuki et al. 2013; Yubuki et al. 2016; Hamann et al. 2017) and large vesicular endosome-like structures observed in *S. salmonicida* and *S. vortens* and the more distantly related Parabasalia member, *T. foetus*, specific and complete remodelling of endosomes has occurred in the *Giardia* genus. *T. foetus*, except for the presence of endosome-like vesicles, presents digestive vacuoles and a stacked Golgi apparatus. Coated vesicles, likely CCVs, are observed near the *T. foetus* Golgi apparatus and the PM. In our analysis, we could not confirm fluid phase material uptake through the cytostome present in *Spironucleus sp* (Sterud and Poynton 2002). As noted, dextran accumulated in spherical vesicles of different dimensions and unknown origin, similar to endosomes. Figures 10A and B summarize the results of our comparative analysis and highlight the unique endocytic system in *Giardia* where, unlike related species and other excavates, PEC-mediated uptake is restricted to the dorsal side of the cell (Ebneter and Hehl 2014; Zumthor et al. 2016) while the ventral side is deputed to attachment to host structures. Interestingly, endosome and lysosome tubulation has been documented in macrophages (Hipolito et al. 2018; Suresh et al. 2020) and are linked with different physiological states of the organelles and subsequent function in the cell – like prompting the cell for phagocytosis. This does permit the hypothesis for the different kinds of PECs present in *G. lamblia* also representing different stages in organelle maturation and or active role at a given time. In line with this, we provide evidence for different stochiometric association of CHC foci with different kinds of PECs. Naturally, only further biochemical dissection of *Giardia* endocytic pathway will help clarify the matter.

### *G. lamblia* possesses a highly divergent clathrin heavy chain and a newly acquired clathrin light chain analogue

We performed an in-depth search for CHC homologs within excavates and other key eukaryotic groups. We found that CHC is conserved in all of these organisms, underlining the vital role of CHC in eukaryotic organisms. The *s*equence divergence of the giardial CHC protein is reflected in an overall decrease in the number of α-helical domains which are essential for the formation of the triskelion leg, and hence necessary for coat assembly (Kirchhausen et al. 2014). Thus, the reduction in α-helical domains during *Gl*CHC evolution, may have led to a lower propensity of *Gl*CHC forming triskelion assemblies and membrane coats. So far, none of the many attempted methods to detect GlCHC in association with small vesicles have been able to show anything other than an exclusive focal localization at PVs/PEC membrane interfaces (Zumthor et al. 2016)). Also, the *Gl*CHC protein does not contain the C-terminal uncoating motif “QLMLT” nor is this motif present in the CHC homologs of any diplomonad. In fact, this motif appears to be only present in Metazoa and in the closely related Filastera and Choanoflagellata (King et al. 2008; Fairclough et al. 2013; Suga et al. 2013) despite the documented ability to form and uncoat *bona fide* CCVs in some protozoa (Link et al. 2021). In Fungi only a partial “L(M)TL”motif was identified. We could not detect a conserved uncoating motif in CHC sequences of members of the Archaeplastida, Amoebozoa or SAR supergroups (supplementary figure 7). Taken together, this data indicates the uncoating QLMLT motif is apparently specific to and likely and invention of the Holozoa lineage. This observation points to as yet uncharacterised uncoating mechanisms are present in other lineages. For example, clathrin mediated endocytosis is essential in the parasitic protist *Trypanosoma brucei* and CCVs have been documented in this organism (Morgan et al. 2001; Allen et al. 2003; Adung’a et al. 2013; Link et al. 2021). Clathrin and other coat proteins associated with CCVs need to be recycled. While HSC70 is documented in *T. brucei* and likely involved in clathrin uncoating (Rapoport et al. 2008), no *bona fide* uncoating motif has been documented (Adung’a et al. 2013; Manna et al. 2017; Link et al. 2021).

In contrast to *Gl*CHC, evolution of the previously identified putative *Gl*CLC/*Gl*4259 protein, presents a different and surprising natural history. This protein was identified as the strongest interactor of GlCHC (Zumthor et al. 2016) and is present in all sequenced *Giardia* lineages. *Gl*CLC/*Gl*4259 has no measurable sequence conservation but a high degree of structural similarity to *bona fide* CLCs, warranting its proposed renaming to *Gl*ACLC. Aside from the *Giardia* genus, we were unable to identify homologues for *Gl*ACLC in any other eukaryotic taxa, nor could we find any orthologues of CLC in any available Fornicata genome/transcriptome sequence, suggesting that the last Fornicata common ancestor (LFCA) lacked a canonical CLC. Taken together, the available data is currently insufficient to decide between two mutually exclusive evolutionary scenarios: 1) secondary loss of a canonical CLC in the last fornicate common ancestor, with acquisition of a structurally and functionally related *GlACLC*, or 2) massive sequence divergence of the original CLC due to significant changes in function particularly in Giardia where originally dynamic membrane coating machinery has evolved to become a static structural element supporting interfaces between plasma membrane and the endocytic system. The discovery of a strong interactor of CHC in the closely related *S. salmonicida Ss*11905, with structural similarity to *GlACLC* as well as to bona fide CLCs is consistent with both scenarios. Notably, this protein neither retrieves *GlACLC* nor CLCs in BLAST searches, leaving no evidence of direct homology. By contrast, robust predictions for CLC homologues were made for members of the Preaxostyla, Discoba and Parabasalia lineages. Nonetheless, other lineages appear also to have lost a bona fide CLC, like *C. parvum* and *T. thermophila* (Woo et al. 2015; Richardson and Dacks 2022), but perhaps similar investigations to ours of CHC may identify CLC analogues/divergent homologues. Taken together, this data suggests that the constraints on the CHC primary structure are higher than on CLC even after massive changes in clathrin coat function with demonstrated complete losses in some protists. Members of the *Giardia* genus as well as *S. salmonicida* have no identifiable *bona fide* CLC, yet, at least the giardial *Gl*ACLC has retained its function as a CHC interacting partner.

Our data provide a robust understanding of *Giardia, Hexamitidae* members, and *Tritrichomonas foetus* endocytic pathway organellar ultrastructure. Contrary to *Spironucleus* or *Tritrichomonas* and other excavates, *Giardia* underwent a complete remodelling of its endocytic machinery. Our analysis revealed its organelles to be polymorphic in nature, justifying the proposed name change to peripheral endocytic compartments. Our analysis of *G*CHC sequences highlights its divergence which is likely due to a massive reorganization of the endocytic pathway in these species, whilst origin and evolution of CLC structural and to some extent functional homologs in *Giardia* (*GlACLC*) and in certain *Hexamitidae* members (*S. salmonicida* and *Trepomonas* sp. PC1) remains uncertain.

## MATERIALS AND METHODS

### Cell culture and transfection

*Giardia intestinalis* strain WB (clone C6; ATCC catalog number 50803) trophozoites were grown using standard methods as described in Morf et. al. (Morf et al. 2010) Episomally-transfected parasites were obtained via electroporation of the circular pPacV-Integ-based plasmid prepared in *E. coli* as described in Zumthor et al. (Zumthor et al. 2016) Transfectants were selected using Puromycin (final conc. 50 μg ml^−1^; InvivoGen). *Spironucleus vortens* and *Spironucleus salmonicida* were cultured as described before (Paull and Matthews 2001; Xu et al. 2014). *S. salmonicida* was transfected using a modified PAC vector and selected with Puromycin (final conc. 50 μg ml^−1^; InvivoGen) (Jerlström-Hultqvist et al. 2012). *Tritrichomonas foetus* was axenically grown also as described (Lealda et al. 1986).

### Construction of expression vectors

*Spironucleus salmonicida* CHC sequence (SS50377_14164) was amplified with the primers <u>ATATTTAATTAAGGCGGATCTATAGTTTCTTGGAATACTAAAATAGGA</u> (forward) and <u>TATGCGGCCGCCACCAGTTATCAGCGGGTGCC</u> (reverse) containing a MluI and a NotI rectrictyion site respectively. The genomic sequence amplified contained a 5’ UTR region of 179bp which encodes a putative promoter. The genomic fragment was inserted in the previously described vector pSpiro-PAC-3xHA-C (Jerlström-Hultqvist et al. 2012).

### Focused Ion Bean Scanning Electron Microscopy (FIB-SEM) of a full *Giardia* trophozoite and image analysis

Wild type *Giardia lamblia* trophozoites were subject to High Pressure Freezing, and processed as established in (Zumthor et al. 2016). Ion milling and Imaging was performed in a Auriga 40 Crossbeam system (Zeiss, Oberkochen, Germany) using the FIBICS Nanopatterning engine (Fibics Inc., Ottawa, Canada) following the aforementioned established protocol. Pixel size was set to 5 nm, obtaining isotropic imaging. Alignment of the dataset was performed resorting to the ImageJ plugin Sift (Schindelin et al. 2012). Image segmentation was done using the semi-autonomous algorithm ilastik (Sommer et al. 2011). The routine of pixel and object classification and used. Algorithm training was performed in a small representative region of the dataset which was then applied to the complete dataset. Imaris (Bitplane AG) was used for three-dimensional rendering and volume measuring.

### Transmission Electron Microscopy analysis of *Giardia lamblia*, *Spironucleus spp*. And *Tritrichomonas foetus* cells and analysis

*G. lamblia*, *S. vortens, S. salmonicida* and *Tritrichomonas foetus* samples were subject to high pressure freezing and processed as we previously established (Gaechter et al. 2008; Zumthor et al. 2016). Samples were imaged in a FEI CM100 Transmission Electron Microscope. Pixel size was assigned to 0.8 nm. Tiles were obtained automatically after determination of focal point. Tiles were aligned with TrakEM2 (Cardona et al. 2012).

### Immunofluorescence Assays

Chemically fixed cells for subcellular recombinant protein localization were prepared as previously described (Konrad et al. 2010). HA-epitope tagged recombinant proteins were detected using a rat-derived monoclonal anti-HA antibody (dilution 1:200, Roche) followed by a secondary anti-rat antibody coupled to AlexaFluor 488 fluorophores (dilution 1:200, Invitrogen). Samples were embedded in Vectashield (VectorLabs) or Prolong Diamond Mounting medium (Invitrogen) containing 4’,6-diamidino-2-phenylindole (DAPI) for nuclear staining.

### Fluid phase marker uptake

Dextran uptake assays were performed as described in (Gaechter et al. 2008; Zumthor et al. 2016) using Dextran 10kDa at 2mg/mL (Invitrogen). Coupled fluorophore was chosen based on image technique chosen. Immunostaining was performed as described above with the exception of using only 0.02% Triton-X100 (Sigma) in 2% BSA (Sigma) for permeabilization, to prevent leakage and loss of Dextran signal. Intensities were calculated with a costume developed macro in Fiji/ImageJ (Schindelin et al. 2012), resorting to WEKA algorithms for segmentation (Arganda-Carreras et al. 2017).

### Laser Scan Confocal Microscopy (LSCM)

Imaging was performed in an inverted Confocal Laser Scanning Microscope Leica SP8 using appropriate parameters. Confocal images were subsequently deconvolved using Huygens Professional (https://svi.nl/Huygens-Professional) and analysed using Fiji/ImageJ (Schindelin et al. 2012).

### Stimulated Emission Depletion (STED) Microscopy

Sample preparation was done as described for LSCM. For imaging, samples were mounted in ProLong Diamond antifade reagent (Thermo Fisher Scientific). Super resolution microscopy was performed on a LSCM SP8 gSTED 3X Leica (Leica Microsystems) using appropriate gating settings. Nuclear labelling was omitted due to possible interference with the STED laser. A pulse depletion laser of 775 nm at 100% strength was used to deplete signal coming from samples using the fluorophore Alexa Fluor 594. Signal from samples containing Alexa Fluor 488 were depleted with the depletion laser line 592 nm at 50% strength. Pinhole was kept at 1 AU. Images were deconvolved using Huygens Professional (https://svi.nl/Huygens-Professional). After deconvolution, signal was segmented following a pixel and object classification routine in ilastik. Thresholding was processed in Fiji/ImageJ (Schindelin et al. 2012) with respective calculation of organelle area.

### Single Molecule Localization Microscopy (SMLM)

Cells were fixed into a coverslip using a cytospin (6 min, 600 g). Samples were then embedded in Vectashield based imaging medium (Olivier et al. 2013). Excess buffer was dried up and samples were sealed. Single Molecule Imaging was performed on a on a Leica SR-GSD 3D microscope (Leica Microsystems) as described in (Mateos et al. 2016) with a cylindrical lenses, in order to image the apical cell region, giving a z-depth of about 800 nm. A minimum of 100 000 events were recorded. Image reconstruction was performed with the ImageJ plugin Thunderstorm (Ovesný et al. 2014). Reconstructed images were segmented following a pixel and object classification routine in ilastik (Sommer et al. 2011; Berg et al. 2019). Thresholding and volume calculation was performed in Imaris (Bitplane AG).

### Native Co-immunoprecipitation of *S. salmonicida* CHC

Co-immunoprecipitation assays on control wild type *S. salmonicida* and transgenic *S. salmonicida* bearing the HA-tagged CHC were processed as previously established (Zumthor et al. 2016) in non-cross-linking conditions agent.

### Protein analysis and sample preparation for mass spectrometry (MS)-based protein identification

SDS-PAGE analysis was performed on 4%-10% polyacrylamide gels under reducing conditions. Blotting was done as described in (Konrad et al. 2010) using primary rat-derived anti-HA antibody (dilution 1:500, Roche) followed by anti-rat (dilution 1:2000; Southern Biotech) antibody coupled to horseradish peroxidase. Gels for mass spectroscopy (MS) analysis were stained with Instant blue (Expedeon) and de-stained with ultrapure water. MS-based protein identification was performed as previously reported (Zumthor et al. 2016).

### In silico co-immunoprecipitation dataset analysis

The co-IP datasets derived from transgenic cells expressing epitope-tagged “baits” as affinity handles were filtered using dedicated control co-IP datasets generated from non-transgenic wild-type parasites to identify candidate interaction partners unique to bait-specific datasets. This was done using Scaffold4 (http://www.proteomesoftware.com/products/scaffold/). Unless otherwise indicated, bait-derived co-IP data was filtered using high stringency parameters (Exclusive Spectrum Counts at 95-2-95, 0% FDR) and manually curated to rank putative interaction partners in a semi-quantitative fashion using ESCs as a proxy for relative abundance. Only proteins with more than 10 hits were considered. Proteins in both datasets were only considered if present 3-fold in the transgenic line versus the control. *In silico* analysis of hypothetical proteins was mainly carried out using BLASTp for protein homology detection (http://blast.ncbi.nlm.nih.gov/Blast.cgi?PAGE=Proteins) and HHPred (http://toolkit.tuebingen.mpg.de/hhpred) for protein homology detection based on Hidden Markov Model (HMM-HMM) comparisons and a cut-off at e-value < 0.05 was implemented to assign *in silico* annotation to otherwise non-annotated proteins of unknown function (Zimmermann et al. 2017).

Protein structure was modelled with the *ab initio* modelling tool AlphaFold (https://alphafold.ebi.ac.uk/) from Alphabet, powered by Google DeepMind (https://deepmind.com/) deep learning neural network algorithms (Jumper et al. 2021; Tunyasuvunakool et al. 2021). Modelling was done via Google Colab in a Jupyter notebook environment(//colab.research.google.com/github/deepmind/alphafold/blob/main/notebooks/).

TM-align calculation was performed online in the server: https://zhanggroup.org/TM-score/. Pymol (The PyMOL Molecular Graphics System, Version 2.0 Schrödinger, LLC.) was used for protein structure prediction visualisation, superimposing and RMSD calculation using the *cealign* command.

### Data Availability

Access to raw mass spectrometry data is provided through the ProteomeXchange Consortium on the PRIDE platform (Perez-Riverol et al. 2019). Data is freely available using project accession number and project DOI. Project DOI/accession number for datasets derived from bait specific and control co-IP MS analyses are as follows: PXD020201.

### Homologue search and Phylogenetic analysis and tree construction

CHC and CLC sequences were probed among several available genomes and transcriptomes with special focus within the fornicata members. Query protein sequences for CHC and CLC from several pan-eukaryotic representatives were obtained and aligned using MUSCLE v.3.8.31 (Edgar 2004) (Supplementary Table 2). Resulting alignments were used to generate Hidden Markov Models using the hmmbuild option and HHMer searches were made on all available genomes with an e-value cutoff to 0.01 (Eddy 2011). Hits were considered valid if reciprocal BLASTp returned a Homo sapiens homologue with a e-value < 0.05. Transcriptome searches were carried out resorting to tBLASTn searches using the *Homo sapiens* and *Monocercomonoides exilis* respective sequences for CHC or CLC. Once a hit was found it was translated into an amino acid sequence and was considered valid if it pulled a *Homo sapiens* homolog with an e-value < 0.05. All found sequences can be found in supplementary tables 2 to 9. Protein domain searches were performed at the Conservate Domain Database (CDD), through the Pfam database (Lu et al. 2020; Mistry et al. 2021). The interPro and SMART platforms were also used for domain classification (Letunic and Bork 2017; Mitchell et al. 2019). Synonymous vs non-synonymous mutation ratio was calculated with an available online software (http://services.cbu.uib.no/tools/kaks) following maximum likelihood parameters.

### Statistical analysis and further used software

All data was analysed for statistical significance and plotted using Prism 9 (Graphpad, https://www.graphpad.com/scientific-software/prism/) software. Images were composed using Affinity Designer software (https://affinity.serif.com/en-gb/). Video processing was made using Da Vinci Resolve v17.3.

## ACKNOWLEDGMENTS

ABH and CF are funded by Swiss National Foundation grant 31003A-166437 and PR00P3_179813, respectively. Imaging and image analysis were performed with equipment from the Centre of Microscopy and Image Analysis (ZMB) of the University of Zurich. We thank the following members of the ZMB for technical and scientific support: Dr. Jana Döhner, Dr. Moritz Kirchmann, Dr. Dominik Hänni, Dr. José Mateos and Dr. Urs Ziegler. Finally we would like to thank members of the Hehl lab for insightful discussions.

## CONFLICTS OF INTEREST

No conflicts of interest

## AUTHOR CONTRIBUTIONS

RS, ABH and CF designed and curated the study. RS performed all experiments and analysed all ensuing experimental data with the exception of *Spironucleus salmonicida* culturing and uptake experiments performed by AA and SS, and SEM experiments performed by JPZ. SVP, RS and JBD performed molecular phylogeny analyses. RS, ABH and CF wrote and revised the manuscript. All authors read and approved the final manuscript prior to submission.

## Supplementary figures

**Supplementary figure 1 – Rendering of a *G. lamblia* trophozoite scanned with FIB-SEM reveals the cell’s inner ultrastructure.** (A) 3D view of acquired FIB-SEM trophozoite data. (B) Single slice showing inner cellular structures such as cytoskeleton elements at the median body (MB), the ventral disk (VD), the endoplasmic reticulum (ER), mitosomes (m) and peripheral vacuoles (PV), highlighted in the region of interest (ROI). (C) Segmentation of different categories of the dataset: cell volume (138 μm^3^), cytoskeleton, endoplasmic reticulum, peripheral vacuoles, small vesicles and mitosomes. (D) Mitosome volume (N = 14, box-plot) was determined post segmentation at an average volume of 0.001093±0.0005698μm^3^ in a 95% confidence interval between [0.0007643, 0.001422] μm^3^.

**Supplementary figure 2 – Cryo-SEM of freeze-fractured trophozoites reveals varying vacuolar morphology in *Giardia lamblia*.** (A) Overview of cryo-preserved *Giardia* trophozoites subjected to freeze-fracture and SEM imaging. Nuclei (N), Endoplasmic Reticulum (ER), Ventral Disk (VD) and peripheral endocytic compartments (PEC) and plasma membrane (PM) are clearly identifiable. (B and C) Insets showing different PEC morphology: vesicular (asterisk) and tubular (hashtag). Scale bar: (A) 2 μm and (B and C) 500 nm.

**Supplementary figure 3 – TEM investigation of *Giardia lamblia* endocytic and secretory pathway.** (A) Overview of a trophozoite. Different PEC structures, vesicular and tubular are observed, together with small vesicles (SV). The N (nucleus) and ER are also highlighted. (B) Close up on tubular PECs (hashtag). (C) Close up on vesicular PECs (asterisk) and SVs (arrowhead). Scale bars: (A) 2 μm, (B) 1 μm and (C) 500 nm.

**Supplementary figure 4 – TEM investigation of *S. salmonicida* endocytic and secretory pathway.** (A) *S. salmonicida* presents vacuolar formations close to the plasma membrane. Cells also present a prominent endoplasmic reticulum (ER; blue-framed inset). (B) Highlight of vacuolar formations (V) and ER. (C) Second cell displaying an abundance of PV close to its plasma membrane. (D) Highlight of vacuoles (V) and the prominent ER that connects to the plasma membrane (asterisk). (E) *S. salmonicida* PVs average a diameter of 205±62.6 nm (N=114) in a 95% confidence interval of [193;217]. (F) *S. vortens* peripheral vacuoles are larger than *S. salmonicida* vacuoles in a statistically significant manner (p-value < 0.0001). Diameters were manually determined. Scale bars: (A and C) 2 μm and (B and D) 500 nm.

**Supplementary figure 5 – TEM investigation of *T. foetus* Golgi vesicles**. (A) More than one Golgi apparatus (G) can be found per cell. These organelles resemble canonical stacked Golgi releasing small coated vesicles. (B) These vesicles average a diameter of 58.4±13.1 nm (N=128) in a 95% confidence interval of [56.1;60.7] nm. Scale bar: (A) 500 nm.

**Supplementary figure 6 – Pan-Eukaryotic prediction of clathrin heavy chain protein domains.** Pfam analysis of predicted protein domains for several clathrin heavy chain proteins sequences from the following species: *Giardia lamblia, Spironucleus vortens, Spironucleus salmonicida, Trepomonas sp., Hexamita inflata, Dysnectes brevis, Kipferlia bialata, Carpediomonas membranifera, Aduncisulcus paluster, Chilomastix cuspidata, Trypanosoma brucei, Naegleria gruberi, Tritrichomonas foetus, Monocercomonoides exilis, Tetrahymena thermophila, Hemimastix kukwestjiik, Chlamydomonas reinhardtii, Dyctiostilium discoideum, Saccharomyces cerevisiae, Caenorhabditis elegans, Homo sapiens, Salpingoeca rosetta, Capsospora owczarzaki* and *Monosiga brevicollis*. A general decrease in domain complexity is observed in excavates compared with higher eukaryotes. CLOs: Carpediomonas-like organisms. Diplom: Diplomonada.

**Supplementary figure 7 – The QLMLT motif is exclusive to Holozoa.** Alignment of the C-terminii of CHC sequences from selected Opisthokonta, Archaeplastida, Amoebozoa and SAR species highlights the present of the QLMLT uncoating motif only in Holozoa supergroup. The positioning of the QLMLT is highlighted in blue.

**Supplemental figure 8 – Calculation of *Giardia* ACLC synonymous vs non-synonymous mutation ratio (ω = ks/kn).** (A) Phylogenetic tree resulting of maximum likelihood analysis of the *Giardia* ACLC sequences. Each node is represented by a number. (B) Overall ω < 1 indicating there is no selective pressure on *Giardia* ACLC.

**Supplemental figure 9 – *S. salmonicida* CHC (*Ss*CHC) is distributed in the cell cytosol in foci and does interact with a structural form of CLC.** (A) *Ss*CHC was tagged C-terminally with three HA tags. It is found throughout the cell cytosol. Signal is observed in 88% of the analyzed cells (N = 171). (B) High resolution imaging of *Ss*CHC using confocal imaging reveals CHC foci. (C) Distribution of the 171 proteins found in higher abundance in the *Ss*CHC native co-IP dataset. (D) Native co-IP of tagged *Ss*CHC reporter reveals interaction with several members of the endocytic pathway such as dynamin or beta-adaptin and calmodulin. (E) *Ab initio in silico* protein modelling with AlphaFold of *Ss*11905, *Gl*ACLC, T*b*CLC and *Hs*CLC. TM-align and RMSD scores for predicted structures of *Giardia* ACLC, *Trypanosoma brucei* CLC and Ss11905 with respect to *Homo sapiens* CLC show overall structural conservation with respect to a *bona fide* CLC. Scale bars: (A) 20 μm. (B) 5 μm.

**Supplementary figure 10 – *Trepomonas* sp. PC1 also harbours a putative CLC analogue.** *Ab initio* protein modelling of TPC1_16039, orthologous to *Ss*11905 in combination with *Ss*11905, *Gl*ACLC, *Tb*CLC and *Hs*CLC. RMSD and TM-align cores show overall structural conservation with respect to a *bona fide* CLC.

## Supplementary Videos

**Supplementary Video 1 –** Three-dimensional rendering of endocytic compartments in *G. lamblia* derived from FIB-SEM sectioning and imaging. Scale bar 1 μm.

**Supplementary Video 2 –** Comparison between confocal and STED imaging of *Giardia* PECs. Scale bar: 3 μm.

**Supplementary Video 3 –** Tri-dimensional reconstruction of PV/PECs from STORM data.

**Supplementary Video 4 –** Tri-dimensional confocal imaging of *S. vortens* with Dextran-Texas Red. Both peripheral and near-nuclear endosome-like vacuoles are observed.

**Supplementary Video 5 -** Tri-dimensional STED imaging of *S. vortens* with Dextran Alexa Fluor 594 reveals endosome-like vacuoles in greater detail.

**Supplementary Video 6 -** Tri-dimensional STED imaging of *T. foetus* with Dextran Alexa Fluor 594 reveals endosome-like vacuoles in greater detail.

**Supplemental Video 7 –** Tri-dimensional high resolution confocal imaging and representation of *Ss*CHC-3xHA foci in the cell cytoplasm.

## Supplementary Tables

**Supplementary Table 1-** PECs volume comparison as calculated in FIB-SEM and STORM experiments

## Supplementary Tables 2-9 (one file)

**Supplementary Table 2** - Queries used for CHC HHM profile building

**Supplementary Table 3** - Results from Pan-Eukaryotic search of CHC homologues in available Proteomes

**Supplementary Table 5** - Queries used for CLC HHM profile building

**Supplementary Table 6** - Results for GlCLC search in available Giardia genomes

**Supplementary Table 7** - Results for bona fide CLC present in other genomes/transcriptomes Supplementary Table 8 - Comparison of GlCLC with bona fide CLC

**Supplementary Table 9** - Ten best hits from HHPred

**Supplementary Table 10** - Search for domains from Clathrin heavy chain super family repeats

**Supplementary Table 11** - Modelled sequences for CLC and alike

**Supplementary Table 12** - SsCHC co-IP results

## Notes

http://www.ebi.ac.uk/pride/archive/projects/PXD020201

### Competing Interest Statement

The authors have declared no competing interest.

### Summary of Updates

One of the co-authors' affiliations had to be updated. Email addresses for all authors except the corresponding author, were removed.

