## supplementary figures 1-10 for "Combined nanometric and phylogenetic analysis of unique endocytic compartments in Giardia lamblia sheds light on the evolution of endocytosis in Fornicata": SFigure8_evolutionGiardiaCLC__20210829.pdf

Supplementary Figure 8

A

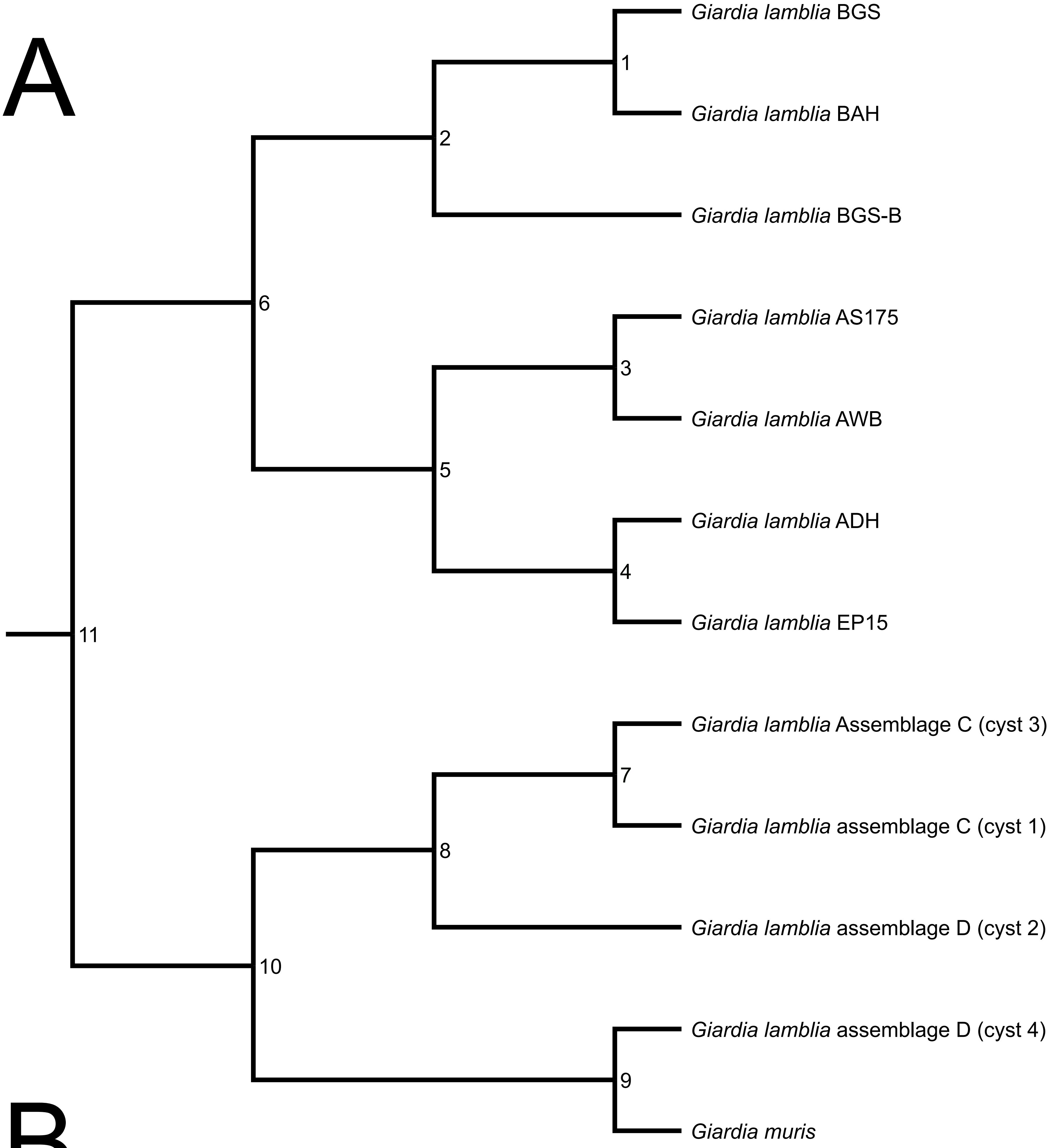


B

| Node# | Ka/Ks Branch1 | Ka Branch1 | Ks Branch1 | Ka/Ks Branch2 | Ka Branch2 | Ks Branch2 |
|---|---|---|---|---|---|---|
| 1 | 0 | 0 | 1.00E-10 | 1.2214 | 0.00389907 | 0.00319237 |
| 2 | 0 | 0 | 1.00E-10 | 0 | 0 | 1.00E-10 |
| 3 | 5.768 | 0.00576798 | 1.00E-10 | 0.6311 | 0.00851468 | 0.01349214 |
| 4 | 1.3124 | 0.02039667 | 0.01554157 | 0.7171 | 0.05336926 | 0.0744284 |
| 5 | 1.0865 | 0.01310214 | 0.01205889 | 1.0467 | 0.01439433 | 0.01375237 |
| 6 | 1.9428 | 0.06120556 | 0.03150349 | 1.2956 | 0.08488583 | 0.06551726 |
| 7 | 0 | 0 | 1.00E-10 | 1.2666 | 0.00126655 | 1.00E-10 |
| 8 | 0.4003 | 0.02242913 | 0.05603452 | 0.8672 | 0.06255149 | 0.07213158 |
| 9 | 0.04253727 | 0.0205944 | 0.4841 | 0.773 | 0.5227 | 0.6762 |
| 10 | 0.0583616 | 0.01207226 | 0.2069 | 0.03765873 | 0.01532114 | 0.4068 |
| 11 | 0.07482582 | 0.04253048 | 0.5684 | 0.05383007 | 0.04210393 | 0.7822 |
