## supplementary figures 1-10 for "Combined nanometric and phylogenetic analysis of unique endocytic compartments in Giardia lamblia sheds light on the evolution of endocytosis in Fornicata": SFigure9_Ss-coIP__prot_comparison_20220326.pdf

Supplemental Figure 9

A

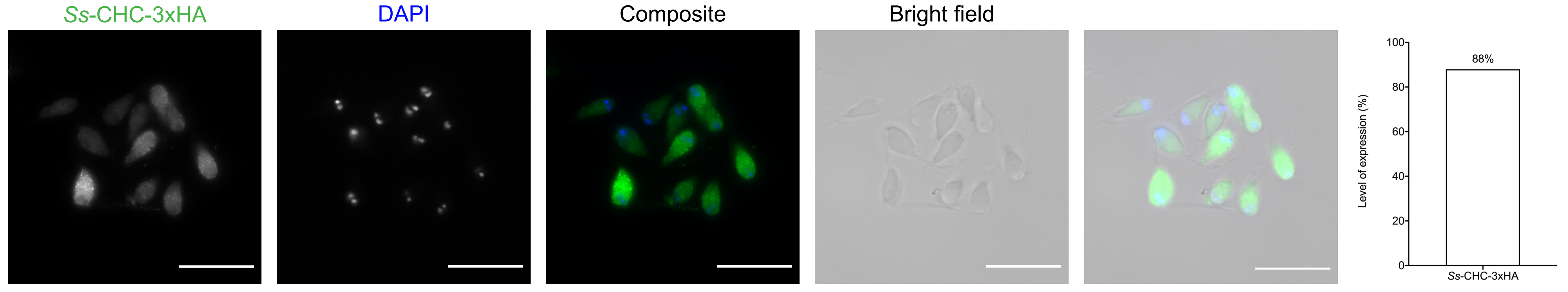


B

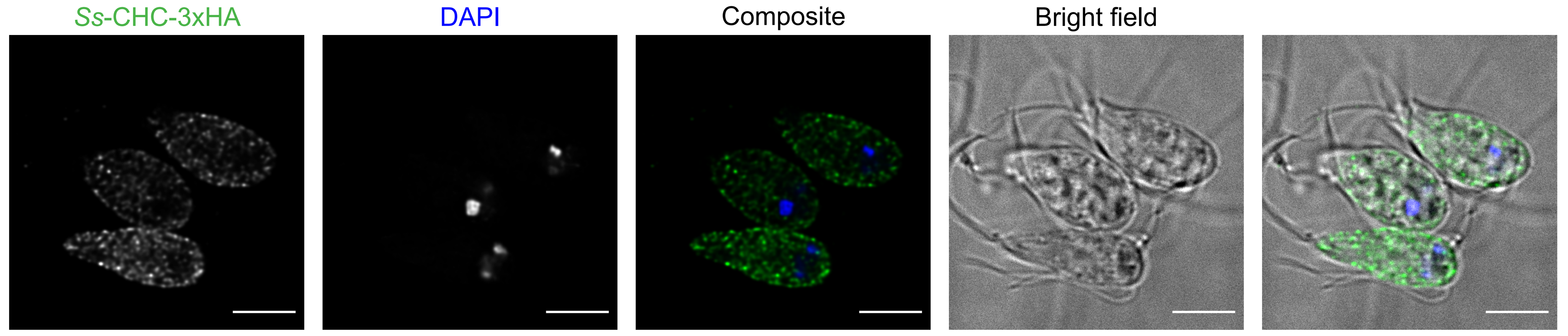


C

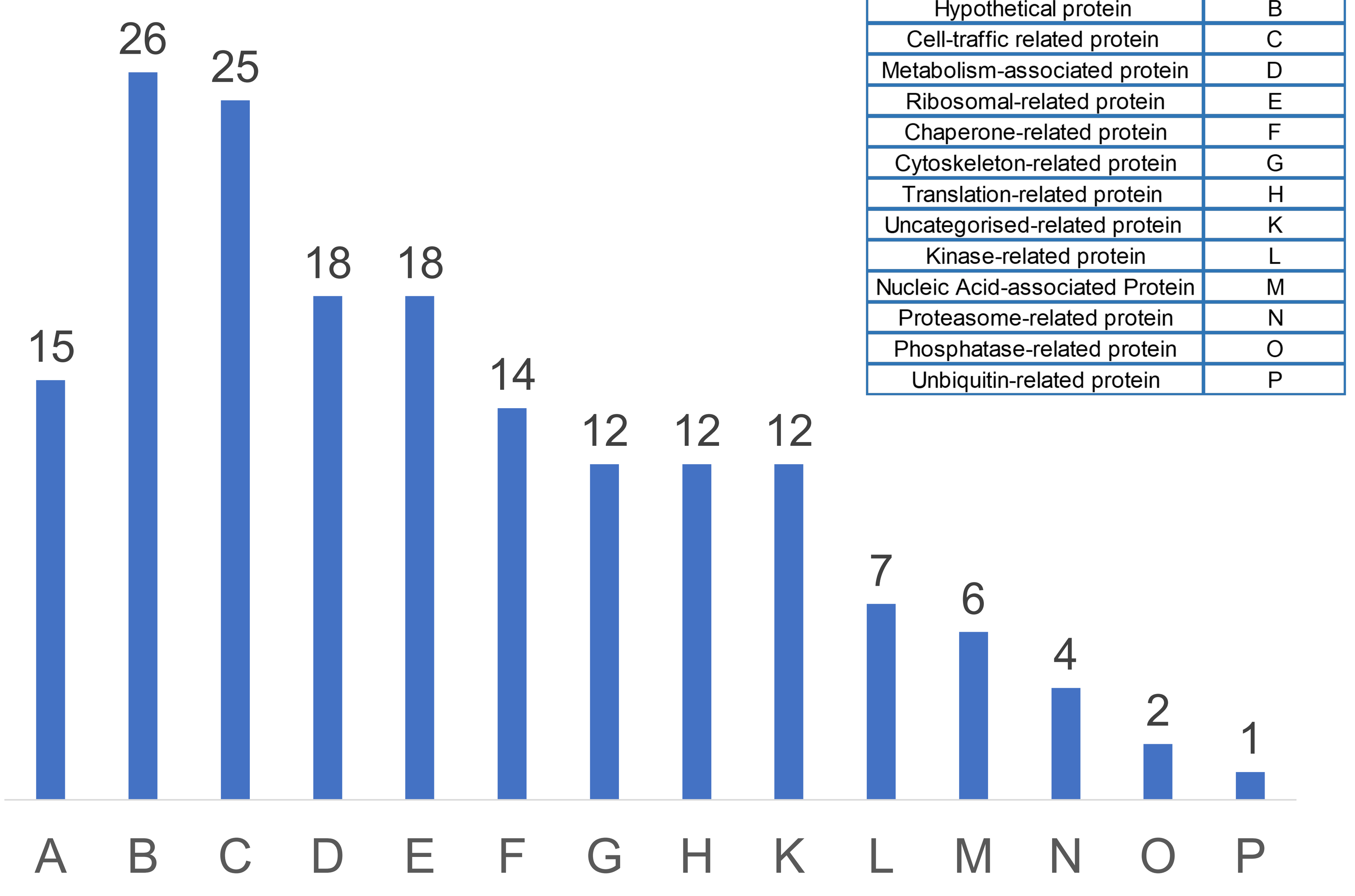


| Protein Group | Graph code |
|---|---|
| Endocytic/Secretory pathway | A |
| Hypothetical protein | B |
| Cell-traffic related protein | C |
| Metabolism-associated protein | D |
| Ribosomal-related protein | E |
| Chaperone-related protein | F |
| Cytoskeleton-related protein | G |
| Translation-related protein | H |
| Uncategorised-related protein | K |
| Kinase-related protein | L |
| Nucleic Acid-associated Protein | M |
| Proteasome-related protein | N |
| Phosphatase-related protein | O |
| Unbiquitin-related protein | P |

D

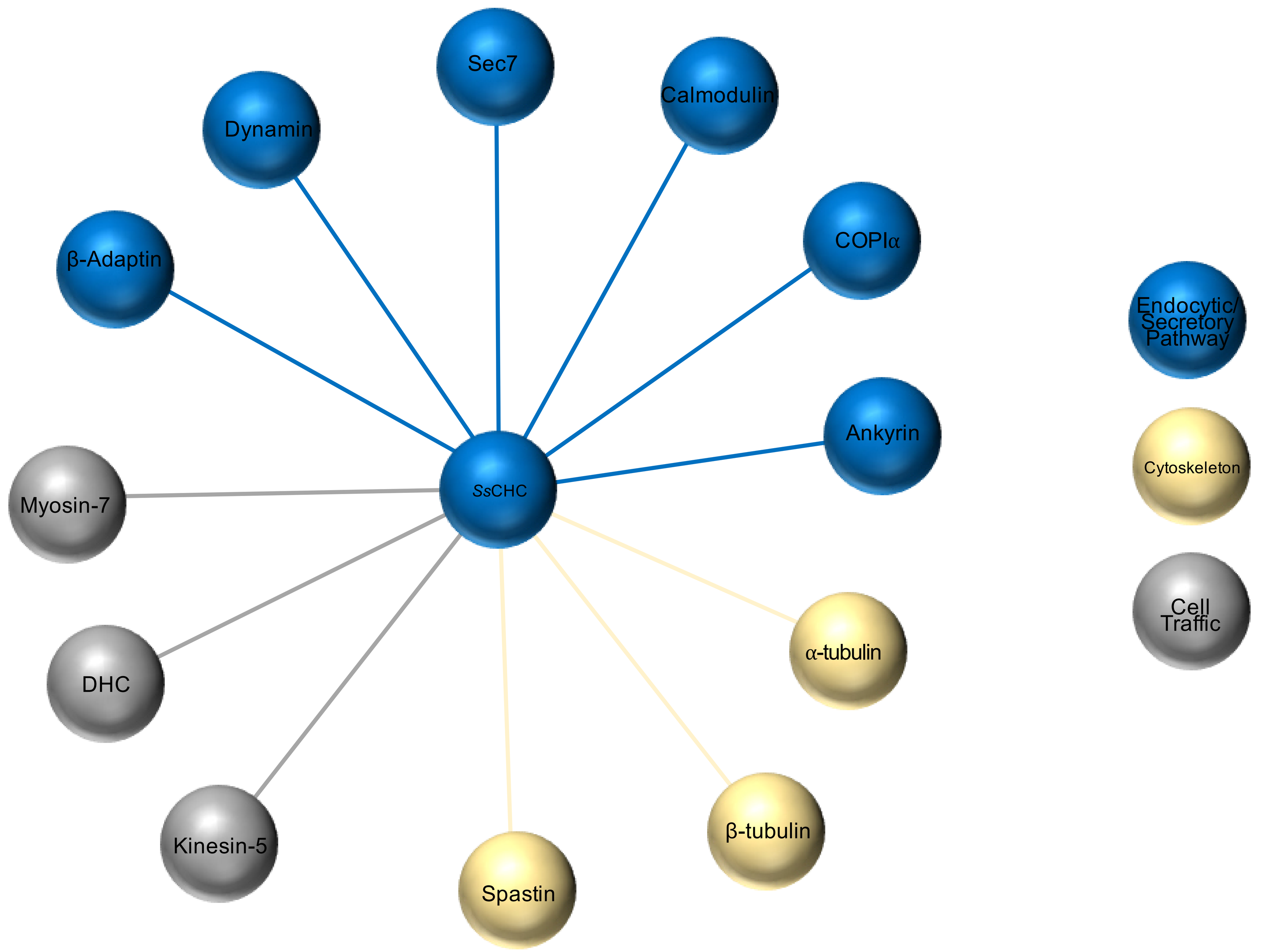


E

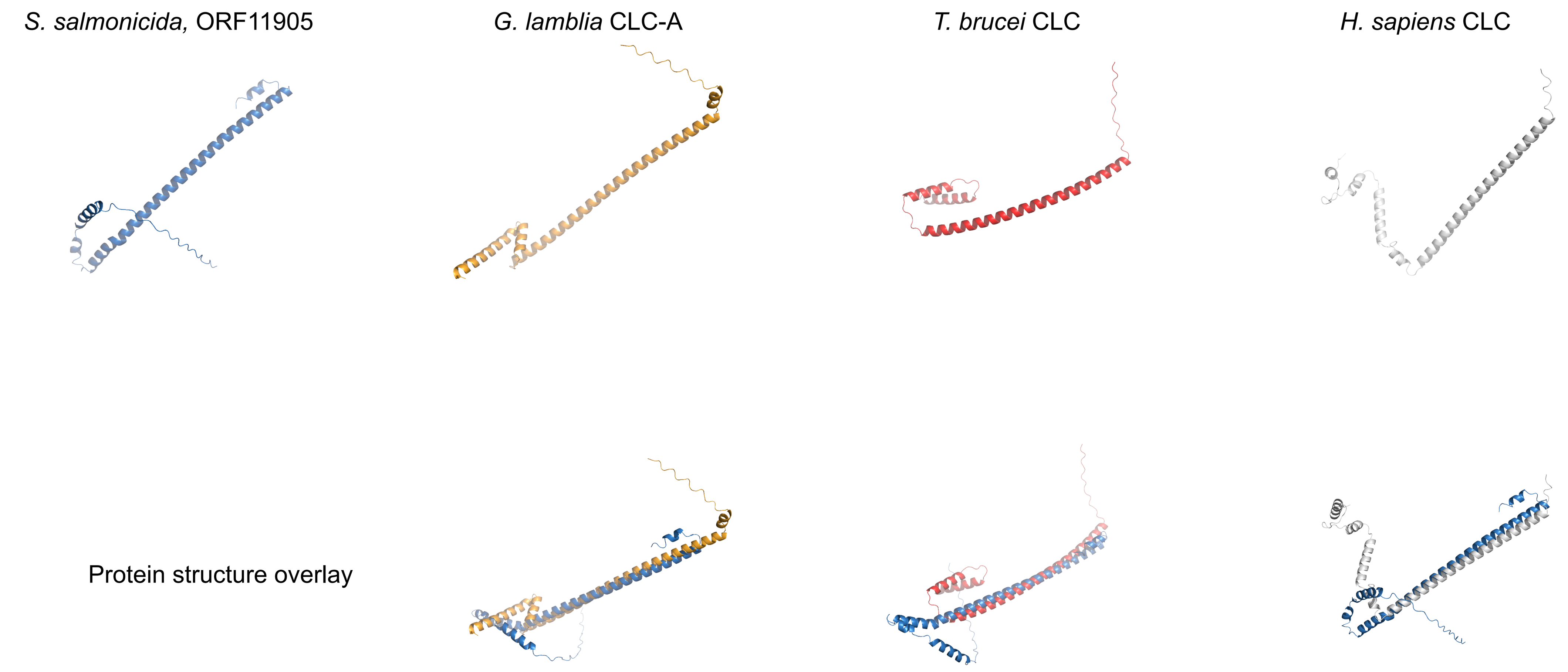


| Specie/TM-align | *H. sapiens* | *T. brucei* | *G. lamblia* | *S. salmonicida* |
|---|---|---|---|---|
| ***H. sapiens*** | 1 | 0.41651 | 0.48174 | 0.43935 |
| ***T. brucei*** | 0.41651 | 1 | 0.47487 | 0.56426 |
| ***G. lamblia*** | 0.48174 | 0.47487 | 1 | 0.55075 |
| ***S. salmonicida*** | 0.43935 | 0.56426 | 0.55075 | 1 |

| Specie/RMSD (Å) | *H. sapiens* | *T. brucei* | *G. lamblia* | *S. salmonicida* |
|---|---|---|---|---|
| ***H. sapiens*** | 0 | 4.58 | 4.58 | 5.32 |
| ***T. brucei*** | 4.58 | 0 | 6.17 | 8.54 |
| ***G. lamblia*** | 4.98 | 6.17 | 0 | 6.62 |
| ***S. salmonicida*** | 5.32 | 8.55 | 6.62 | 0 |
