## supplementary figures 1-10 for "Combined nanometric and phylogenetic analysis of unique endocytic compartments in Giardia lamblia sheds light on the evolution of endocytosis in Fornicata": SFigure10_trepomonas_ACLC_20220306.pdf

Supplemental Figure 10

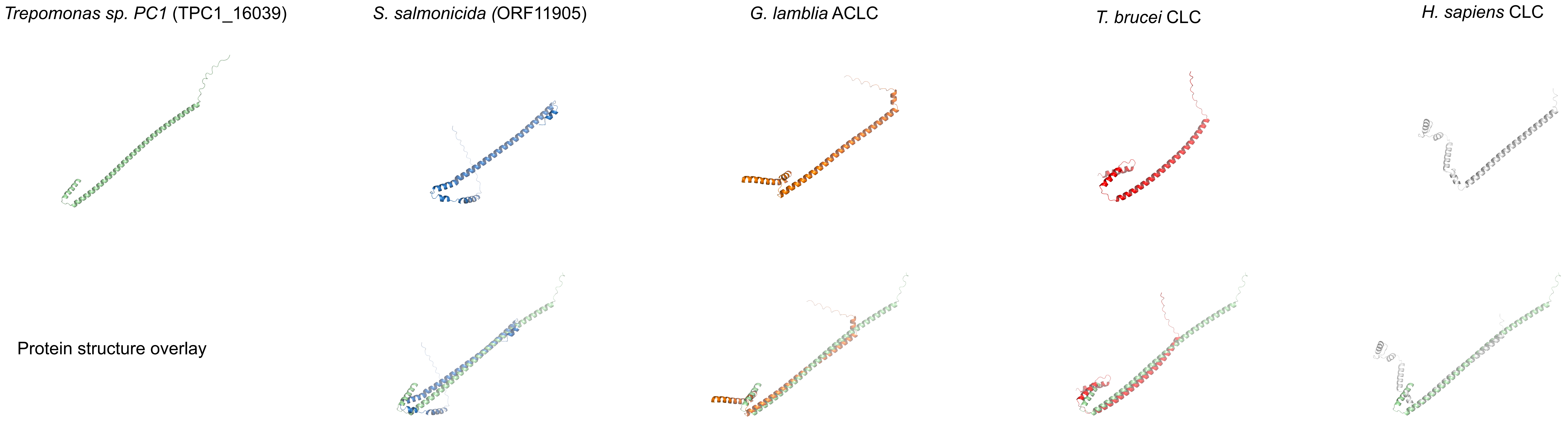


| Specie/TM-align | *H. sapiens* | *T. brucei* | *G. lamblia* | *S. salmonicida* | *Trepomonas sp.* PC1 |
|---|---|---|---|---|---|
| *H. sapiens* | 1 | 0.41651 | 0.48174 | 0.43935 | 0.471 |
| *T. brucei* | 0.41651 | 1 | 0.47487 | 0.56426 | 0.46 |
| *G. lamblia* | 0.48174 | 0.47487 | 1 | 0.55075 | 0.56 |
| *S. salmonicida* | 0.43935 | 0.56426 | 0.55075 | 1 | 0.5 |
| *Trepomonas sp.* PC1 | 0.471 | 0.46 | 0.56 | 0.5 | 1 |

| Specie/RMSD (Å) | *H. sapiens* | *T. brucei* | *G. lamblia* | *S. salmonicida* | *Trepomonas sp.* PC1 |
|---|---|---|---|---|---|
| *H. sapiens* | 0 | 4.58 | 4.58 | 5.32 | 3.1 |
| *T. brucei* | 4.58 | 0 | 6.17 | 8.54 | 4.69 |
| *G. lamblia* | 4.98 | 6.17 | 0 | 6.62 | 3.71 |
| *S. salmonicida* | 5.32 | 8.55 | 6.62 | 0 | 6.45 |
| *Trepomonas sp.* PC1 | 3.1 | 4.69 | 3.71 | 6.45 | 0 |
