## Supplementary figures and images for "Combined nanometric and phylogenetic analysis of unique endocytic compartments in Giardia lamblia sheds light on the evolution of endocytosis in Fornicata"

### SFigure1_FIB_Reconstruction_20220326.pdf

#### Supplemental figure 1

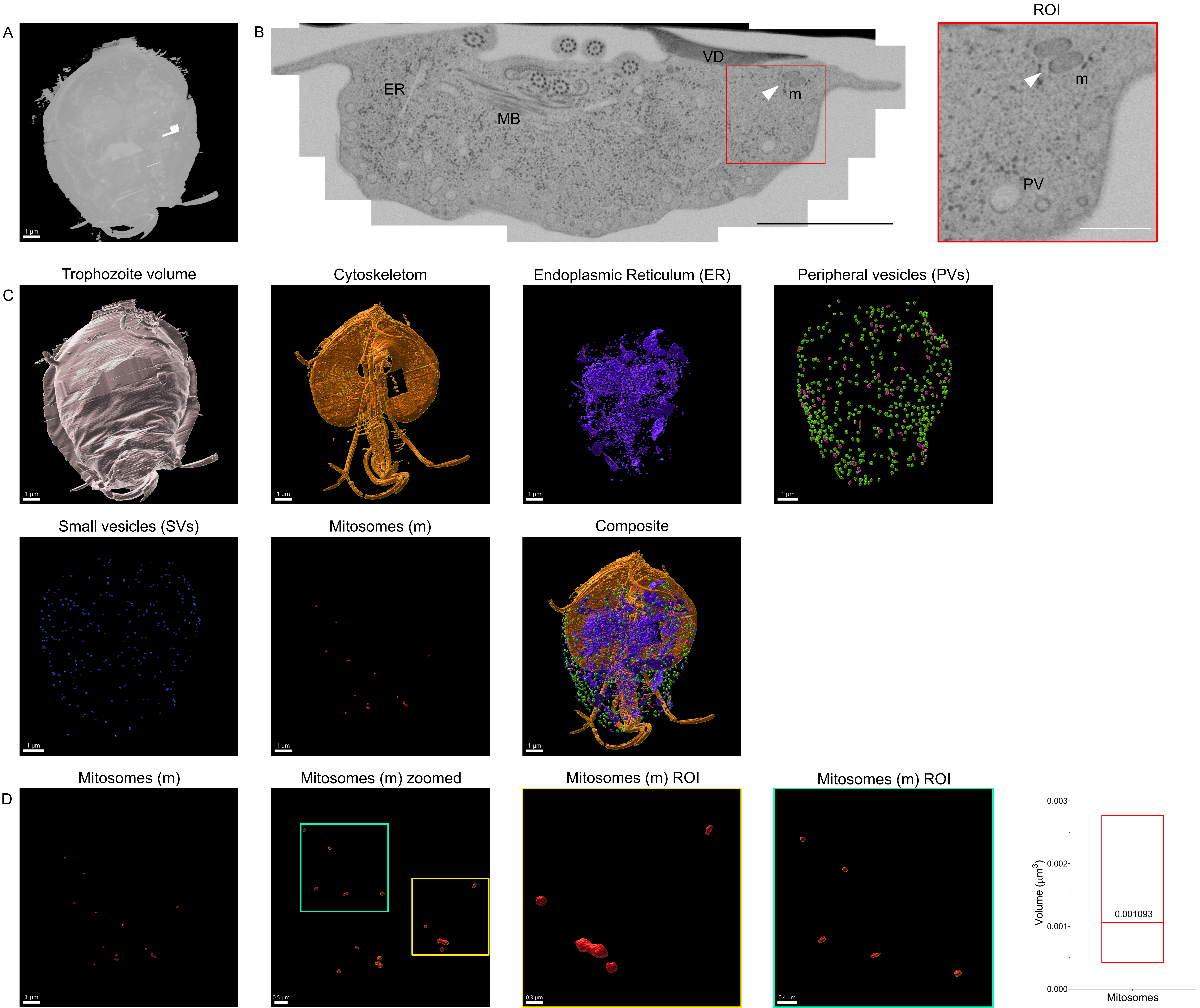

### Sfigure2_Giardia_cryoSEM_20210829.pdf

#### Supplementary Figure 2

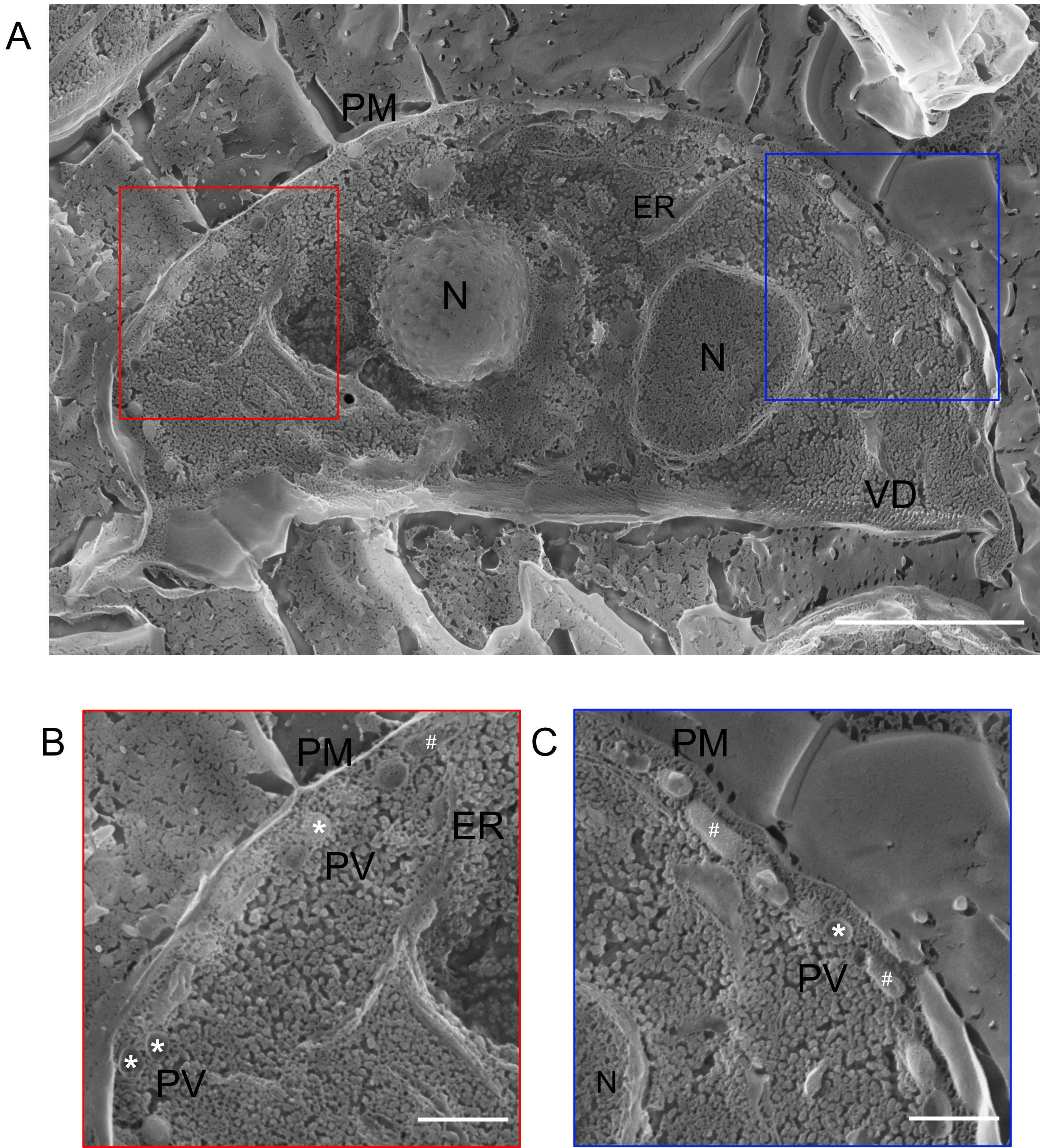

### Sfigure3_WB_TEM__20210829.pdf

Supplementary Figure 3

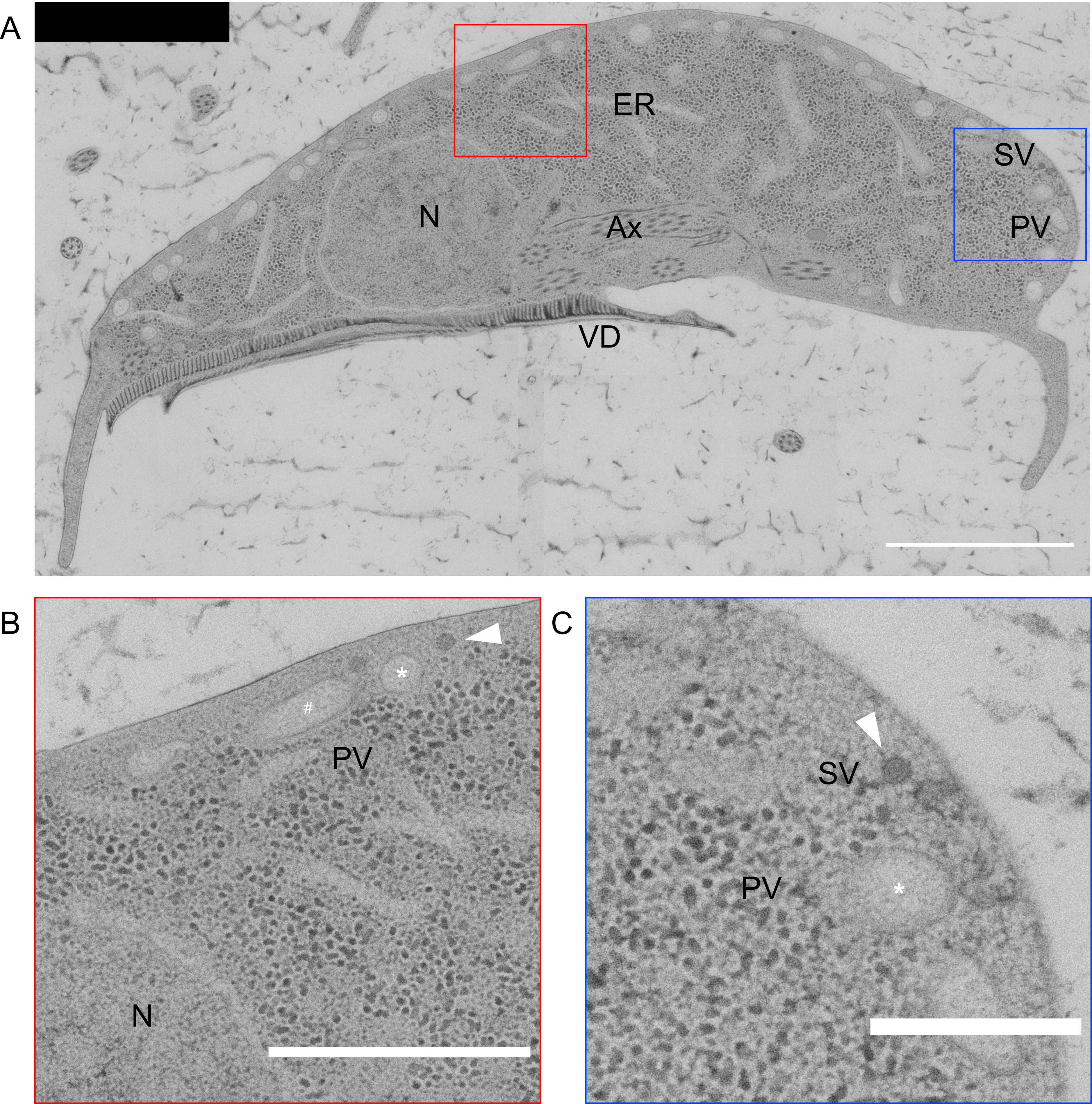

### Sfigure4_Ssalmonicida_TEM__20220326.pdf

#### Supplementary Figure 4

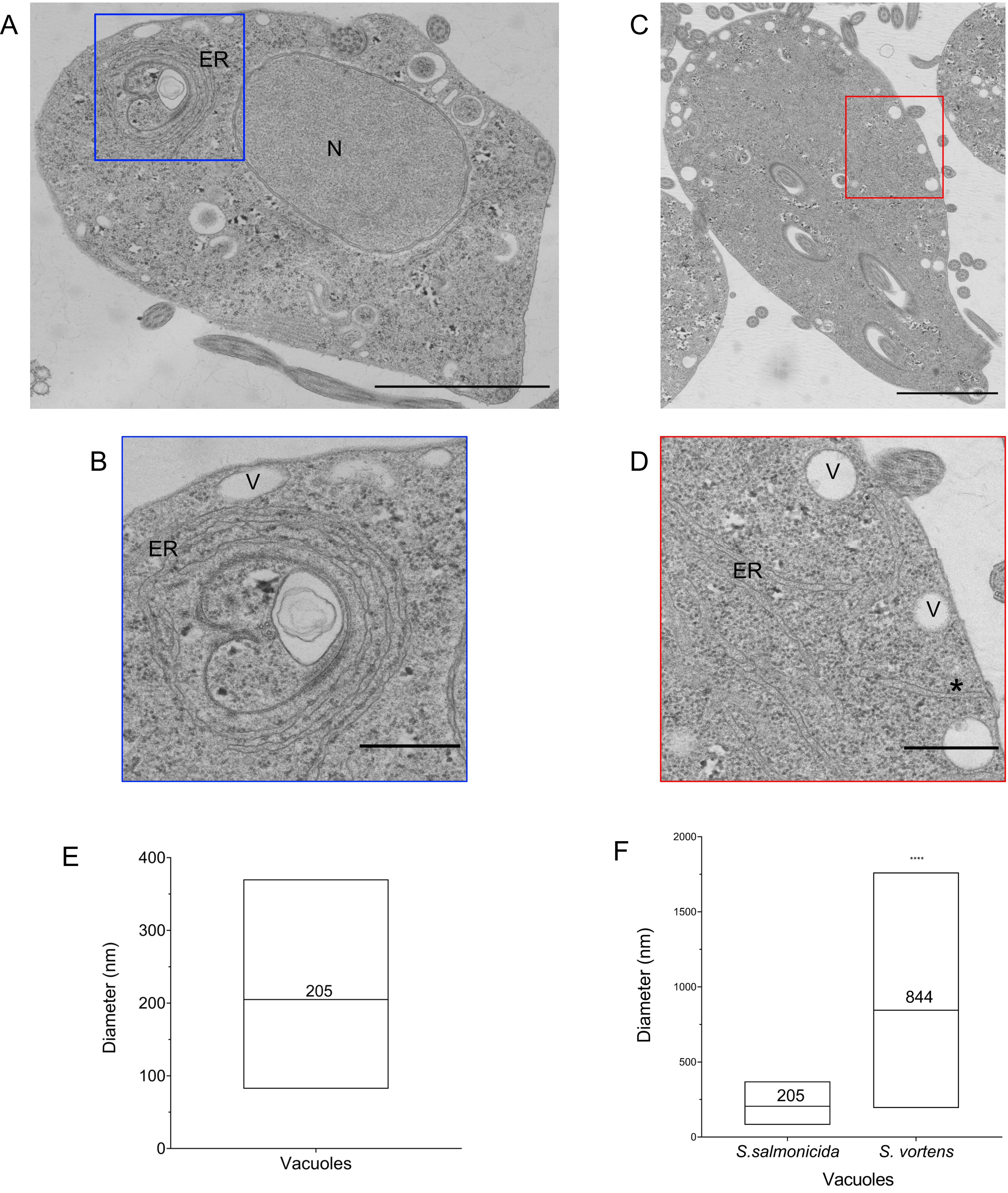

### Sfigure5_Tfoetus_Golgi__20210829.pdf

Supplementary Figure 5

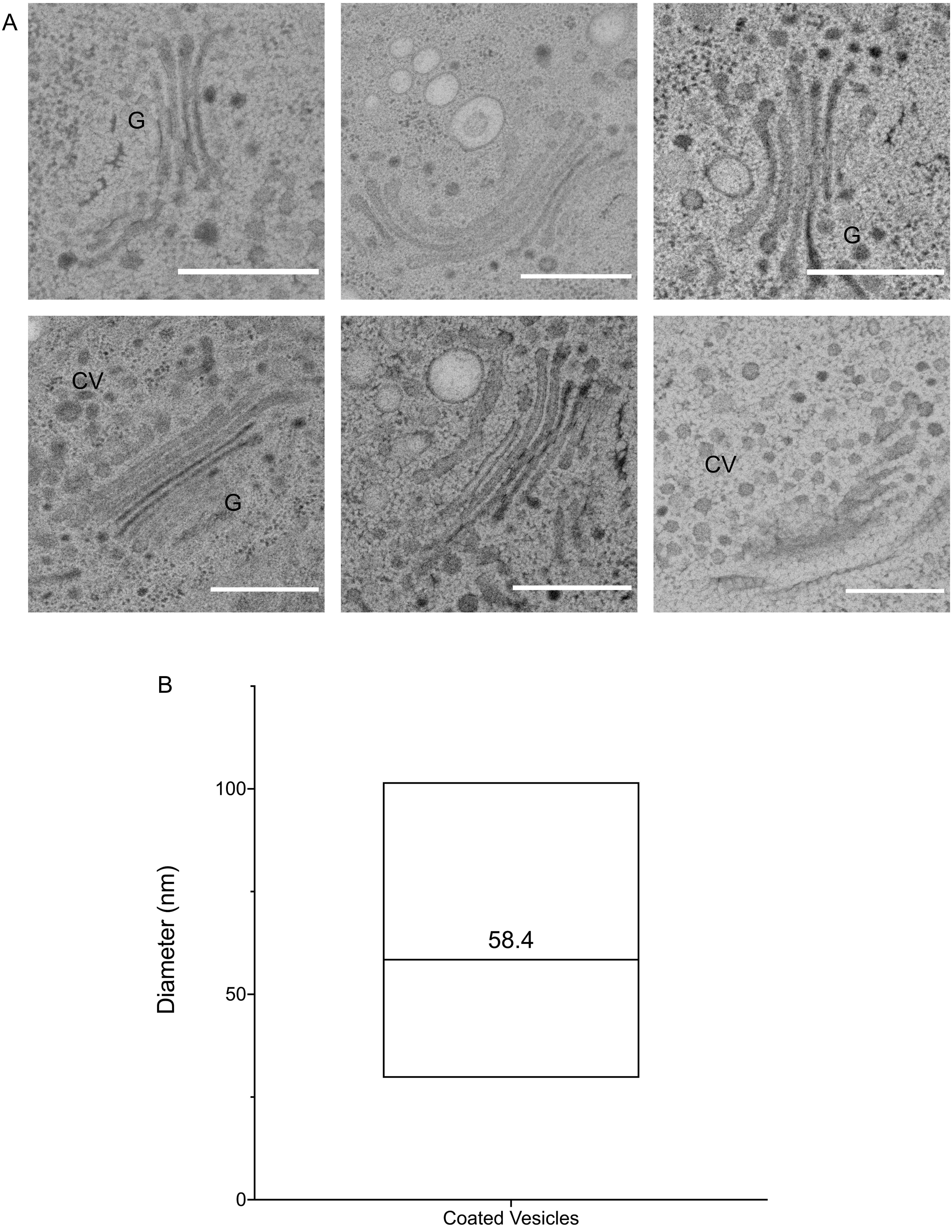

### SFigure6_PanEuk_CHC_prot-domains__20220326.pdf

Supplementary Figure 6

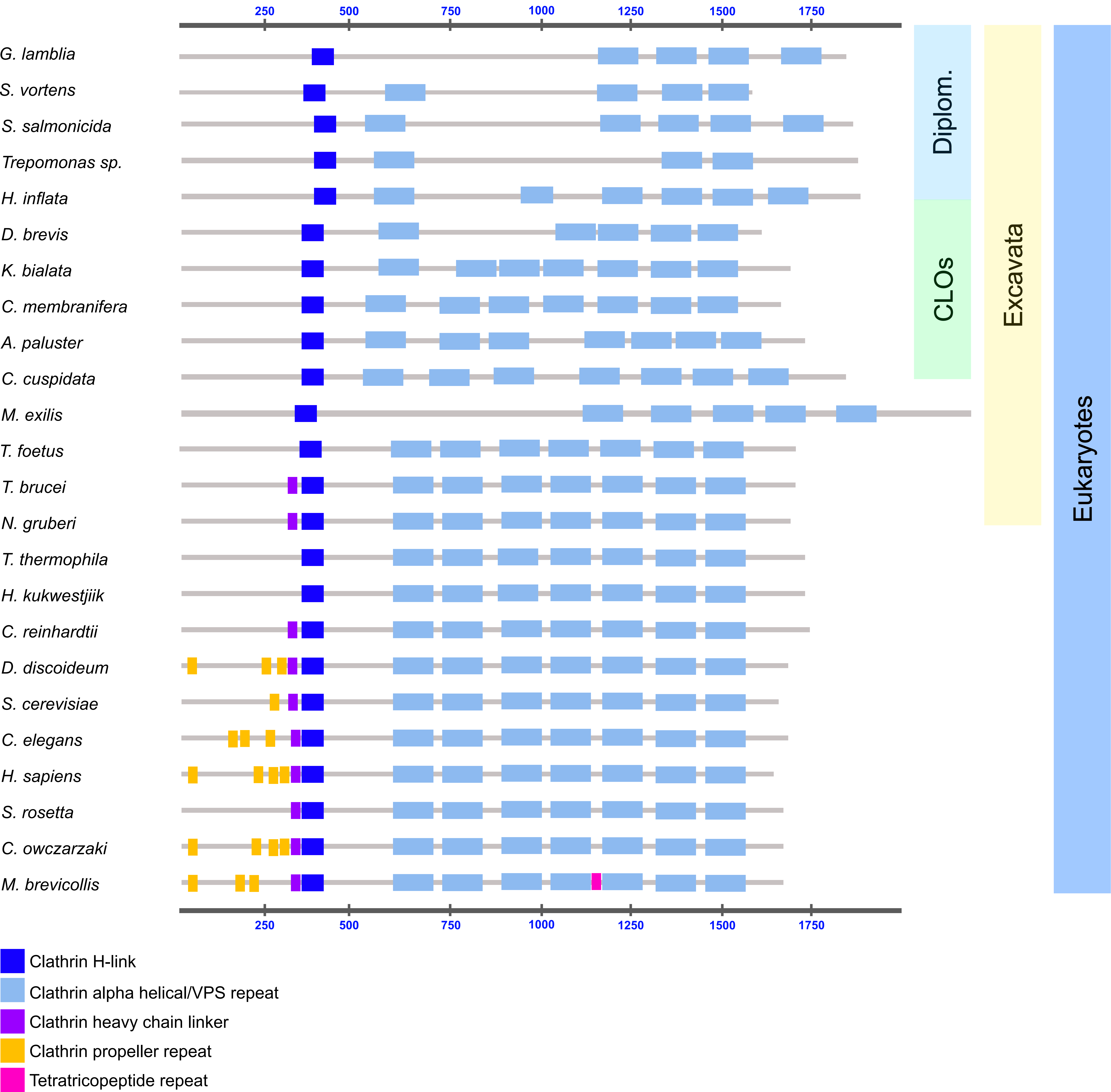

### SFigure7_CHCalignment__20210829.pdf

#### Supplementary Figure 7

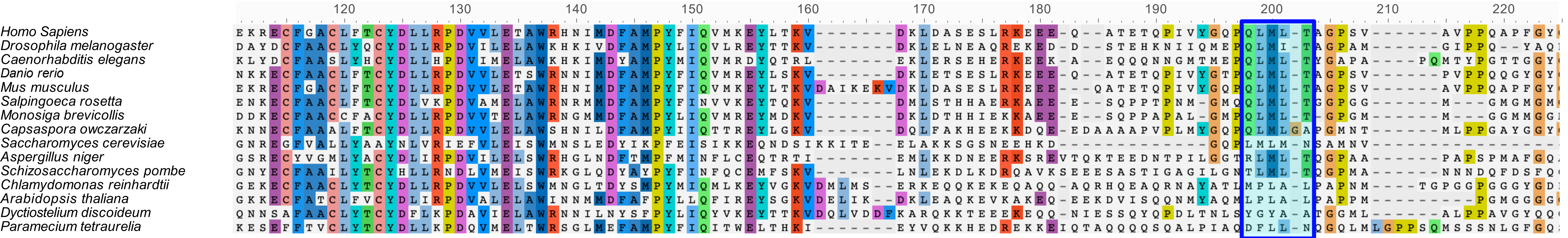
